# *Teosinte Pollen Drive* guides maize diversification and domestication by RNAi

**DOI:** 10.1101/2023.07.12.548689

**Authors:** Benjamin Berube, Evan Ernst, Jonathan Cahn, Benjamin Roche, Cristiane de Santis Alves, Jason Lynn, Armin Scheben, Adam Siepel, Jeffrey Ross-Ibarra, Jerry Kermicle, Rob Martienssen

## Abstract

Meiotic drivers subvert Mendelian expectations by manipulating reproductive development to bias their own transmission. Chromosomal drive typically functions in asymmetric female meiosis, while gene drive is normally postmeiotic and typically found in males. Using single molecule and single-pollen genome sequencing, we describe *Teosinte Pollen Drive*, an instance of gene drive in hybrids between maize (*Zea mays ssp. mays*) and teosinte *mexicana* (*Zea mays ssp. mexicana*), that depends on RNA interference (RNAi). 22nt small RNAs from a non-coding RNA hairpin in *mexicana* depend on *Dicer-Like 2 (Dcl2)* and target *Teosinte Drive Responder 1 (Tdr1),* which encodes a lipase required for pollen viability. *Dcl2*, *Tdr1*, and the hairpin are in tight pseudolinkage on chromosome 5, but only when transmitted through the male. Introgression of *mexicana* into early cultivated maize is thought to have been critical to its geographical dispersal throughout the Americas, and a tightly linked inversion in *mexicana* spans a major domestication sweep in modern maize. A survey of maize landraces and sympatric populations of teosinte *mexicana* reveals correlated patterns of admixture among unlinked genes required for RNAi on at least 4 chromosomes that are also subject to gene drive in pollen from synthetic hybrids. *Teosinte Pollen Drive* likely played a major role in maize domestication and diversification, and offers an explanation for the widespread abundance of “self” small RNAs in the germlines of plants and animals.

## Introduction

The introduction of novel genetic variation through hybridization is an important evolutionary catalyst^1^, as adaptive introgression in hybrid individuals can increase fitness under new environmental conditions and lead to geographical expansion and diversification^2^. Modern maize, for example, was first domesticated from a close relative of *Zea mays* ssp. *parviglumis* (teosinte *parviglumis*) in the lowlands of southwest Mexico approximately 9000BP, but admixture from a second teosinte, *Zea mays* ssp. *mexicana,* 4000 years later, appears to have catalyzed rapid expansion across the Americas^3^. The combination of divergent genomes, however, also carries inherent risks.

Allelic incompatibilities, deleterious mutations, and differences in gene dosage can all be unmasked following outbreeding^4,5^, resulting in hybrid sterility, inviability, and necrosis^6–9^. The Bateson-Dobzhansky-Muller (BDM) model accounts for such scenarios, proposing that duplicate deleterious mutations can evolve in distinct populations and are only ever made manifest following hybridization. Increasing evidence suggests that at least some of these incompatibilities stem from intragenomic conflict triggered by selfish genetic elements (SGEs)^10–13^.

Meiotic drive depends on selfish elements that actively manipulate reproductive development in order to facilitate their own preferential transmission^14^. Chromosomal drive refers to the manipulation of chromosome segregation during asymmetric female meiosis, as centromeres, heterochromatic knobs, and telomeres exert mechanical advantages that favor their inclusion in the egg cell^15–19^. Examples include *Abnormal 10* (*Ab10*) in both maize and teosinte populations^20,21^. Gene drive on the other hand, occurs preferentially in males and is achieved via disruption of post-meiotic reproductive development resulting in segregation distortion (SD)^22,23^. These systems tend to occur in sperm or haploid spores and involve toxin-antidote (or distorter-responder) pairs in close genetic linkage. Gametes that do not inherit the drive locus are selectively killed, resulting in overrepresentation of the driver^14^. The mouse *t*-complex^24–27^, *Drosophila Segregation Distorter* (*SD*) complex^28,29^, and *S. pombe/kombucha wtf* spore killers^30,31^ are all autosomal drivers that selectively kill competing wild-type gametes in heterozygotes.

Meiotic drive has significant evolutionary consequences^14,15^. Under simple assumptions, an efficient driver can generally be expected to increase in frequency over time. For X-linked distorters in heterogametic males, this can result in severe sex imbalances and even population extinction^32^.

Because drivers often impose fitness and fertility penalties, tremendous selective pressure is placed on regions of the genome that can evolve suppressors^33^. As a consequence, drive systems undergo recurrent cycles of suppression and counter-suppression^14^. This has a direct impact on patterns of adaptive evolution, and evolutionary conflict in the haploid stage is thought to be a primary reason for unique patterns of germline expression relative to other tissues^34^. Though drive is predicted to be widespread, only a handful of systems have been well-characterized as most drive systems exist in a cryptic state, either through suppression or fixation^14,35^. It is through hybridization with naïve individuals that suppression is lost and drive is once again apparent. The unmasking of cryptic drivers in hybrid individuals likely plays an outsized role in mediating hybrid sterility in plants as well as animals^36^, reinforcing species barriers, and influencing patterns of introgression in hybrid individuals via genetic linkage^37,38^.

Here, we characterize a novel, male-specific SD system in introgression lines between maize (*Z. mays ssp. mays*) and teosinte *mexicana* (*Zea mays ssp. mexicana*) called *Teosinte Pollen Drive* (*TPD*). A pre-meiotic toxin kills pollen grains in a sporophytic manner, and at least two antidotes act as gametophytic suppressors, restoring viability in pollen. We implicate *Dicer-like 2* (*Dcl2*)- dependent small interfering RNAs (siRNAs), as the primary factor mediating pollen killing. These siRNAs are highly abundant in maize pollen and the toxin locus, *Teosinte pollen drive 1* (*Tpd1*), encodes a *mexicana-*specific long non-coding hairpin RNA in close genetic linkage with the centromere of chromosome 5 and a large paracentric inversion. Processing of this hairpin results in specific 22nt siRNAs that target an essential gene *in trans*, resulting in pollen abortion. Co-segregation of a genetically linked hypomorphic (partially functional) *Dcl2* allele suppresses this effect via the reduction of secondary 22nt siRNAs, and is reinforced by a second unlinked antidote (*Tpd2*) on chromosome 6. Single-pollen grain sequencing reveals the biased transmission of several other intervals that also include genes required for RNAi. Survey sequencing of modern and traditional Mexican varieties of maize, and sympatric populations of teosinte indicate that *TPD* has had a substantial impact on patterns of *mexicana* introgression, and on maize dispersal and domestication. Remarkable parallels with endogenous siRNAs in animals suggest that germline RNAi is a general instigator of intragenomic conflict, prompting the idea that “self” small RNAs (those that target genes) can act as pervasive selfish genetic elements.

## Results

### *Teosinte Pollen Drive (TPD)* results in non-Mendelian pollen abortion in Teosinte-Maize hybrids via segregation distortion

Hybridization between maize and teosinte is subject to unilateral cross-incompatibility^39,40^, but pollination of maize by *mexicana* pollen is frequent^41^. Consistently, genome-wide assessments of introgression in sympatric collections have provided evidence for asymmetric gene flow from *mexicana* to maize^41,42^. To further explore the reproductive consequences of hybridization, multiple sympatric collections of *mexicana* were crossed to the Mid-western U.S. dent inbred W22, resulting in variable rates of pollen abortion that typically decreased in subsequent generations. However, a subset of late backcross (BC) lines (hereafter *TPD*) displayed an unusually consistent rate of pollen abortion (75.5% ± 2.48) relative to W22 (6.02% ± 2.95, P < 0.0001, Welch’s t-test) despite normal vegetative and reproductive development (Figure 1a-c; Extended Data Fig. 1a). Interestingly, the pollen abortion phenotype was absent after three rounds of selfing in *TPD* BC8S3 plants (6.40% ± 2.26, P < 0.0001, Welch’s t-test) suggesting that heterozygosity was required (Figure 1d). In reciprocal crosses, pollination of *TPD* ears with W22 pollen resulted in the independent assortment of fertile, semi-sterile and fully male sterile progeny in a 2:1:1 ratio (Figure 1e; Supplementary Table 1). These results indicated the presence of two unlinked loci responsible for pollen survival that were transmitted in a selfish manner to all individuals in the next generation, but only through pollen.

**Figure 1.**
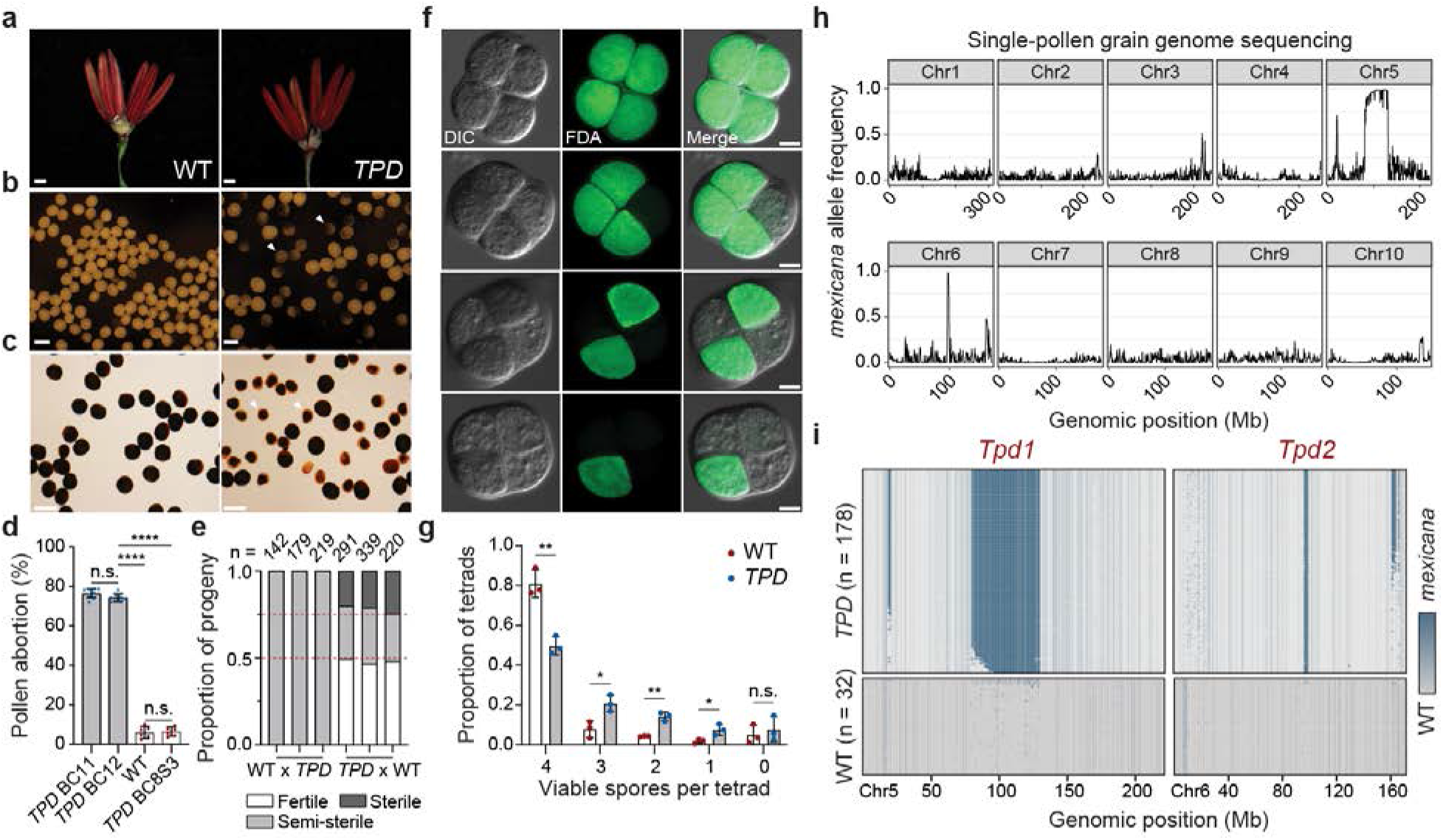
**Single pollen sequencing reveals selfish inheritance in *Teosinte Pollen Drive*. a**, Representative 5mm anther florets from wild-type (WT) and *Teosinte Pollen Drive* (*TPD*) plants. Scale bar = 1mm. **b**, Brightfield imaging of mature pollen grains from WT and *TPD* plants. Arrows denote developmentally arrested pollen grains. Scale bar = 0.1mm. **c**, Iodine potassium iodide (I2KI) viability staining of mature WT and *TPD* pollen grains. Viable pollen grains are plump and darkly stained, whereas arrested pollen grains exhibit reduced diameter and incomplete staining. Scale bar = 0.1mm. **d**, Quantification of pollen abortion rates in *TPD* backcross (BC11,12), WT, and *TPD* self-fertilized (BC8S3) lines. Data are mean ± s.d. (*n* = 6-8). **** p < 0.0001 (Two-tailed *t*-test). **e**, Phenotypic segregation ratios across replicate reciprocal crosses. *n* represents the sample size for each progeny population. The red dashed lines denote a perfect 2:1:1 phenotypic segregation ratio. **f**, Fluorescein diacetate (FDA) viability staining of tetrads from *TPD* plants. Pollen viability is progressively restricted to a single spore following meiosis. Panels show DIC, FDA, and merged images. Scale bar = 50µm. **g,** Viability scoring of *TPD* and WT tetrads shown in f. *TPD* spores exhibit significantly reduced viability at the tetrad stage. n = 3 biological replicates, 952 total tetrads assayed. Data are mean ± s.d. * p < 0.05; ** p < 0.01 (Welch’s t-test). **h**, Single pollen grain genome sequencing. Imputed allele frequencies at *mexicana* markers in a population of 178 mature pollen grains collected from *TPD* plants. **i**, Imputed *mexicana* marker density on chromosomes 5 and 6 for individual pollen grain genome sequences. Multiple *mexicana* haplotypes (blue) are selfishly inherited in viable *TPD* pollen grains (n=178) but not WT (n=32). Values shown (**h**, **i**) are plotted using a 500kb sliding window.

Because this phenotype was observed only in heterozygotes, we reasoned that it stemmed from an incompatibility between the W22 genome and regions of *mexicana* introgression after meiosis, reminiscent of genic drivers that distort patterns of inheritance via selective gamete killing^27,31^.

Consistently, meiotic progression in *TPD* plants was normal until the tetrad stage, following the separation of each haploid complement (Figure 1f). This phenotype, while strictly post-meiotic, appeared to progress gradually, ultimately resulting in arrested pollen grains with a heterogenous overall diameter and varying degrees of starch accumulation (Figure 1c,g).

Genetic mapping revealed that *brittle endosperm 1* (*bt1*) on chromosome 5 and *yellow endosperm 1* (*y1*) on chromosome 6 were linked with the pollen abortion phenotype (Extended Data Fig. 1b,c). Strikingly, backcrosses to *y1*; *bt1* yielded 100% *Bt1* kernels instead of 50%, but only when *TPD* was used as a pollen parent (Extended Data Fig. 1b). The frequency of white kernels (*y1*) was in agreement with recombination estimates (21-22%). This bias was strongly indicative of gene drive, though we could not formally exclude other forms of incompatibility that also result in segregation distortion such as the postzygotic hybrid sterility observed in the Killer-Protector system in rice^43^. To exclude such possibilities, we sequenced the genomes of two homozygous *TPD* lines (BC8S3 and BC5S2) to define 408,031 high-confidence SNPs corresponding to regions of *mexicana* introgression. Next, we sequenced the genomes of individual surviving pollen grains from *TPD* plants, rationalizing that if segregation distortion was occurring in pollen, the causative regions would be overrepresented. We found that several intervals occurred at much higher frequencies than expected after 8 backcrosses (Figure 1h). Notably, introgression intervals on chromosomes 5 and 6 were consistently observed in all surviving pollen (Figure 1i), strongly indicative of post-meiotic gene drive. We designated these loci as *Tpd1* and *Tpd2*, respectively. Thus, non-Mendelian inheritance of pollen abortion was the direct result of SD occurring in pollen. Gametes that could potentially give rise to fertile siblings in maternal progeny were being selectively eliminated in the male germline. As a result, only those pollen grains containing both *Tpd1* and *Tpd2* were contributing to the next generation.

### *Tpd1, Tpd2,* and *Dicer-like2* form a novel toxin-antidote complex

In order to determine the relative contributions of *Tpd1* and *Tpd2* to pollen abortion, we separated the components by maternal transmission into fertile, semi-sterile (“drive”) and fully sterile classes (Figure 2a). Each progeny class had distinct rates of pollen abortion (Figure 2b) and showed significant differences in flowering time (Figure 2c). Fertile segregants were phenotypically wild-type and showed no transmission defects whereas drive plants recapitulated the canonical *TPD* pollen abortion phenotype. In contrast, male reproductive development in sterile plants was developmentally retarded, displaying severely delayed anthesis and reduced overall shed (Figure 2a,c). Consequently, crosses performed with this pollen showed minimal seed set and often failed entirely. We collected pools of plants from the fertile and sterile phenotypic classes (Figure 2d) for bulk segregant analysis (BSA)^44^, and found that *Tpd1* was differentially enriched in sterile plants, while *Tpd2* was enriched in fertile plants (Figure 2e). This indicated that *Tpd1* alone was sufficient to ‘poison’ the male germline, and that this most likely occurred pre-meiotically, as only a single copy of *Tpd1* was required.

**Figure 2.**
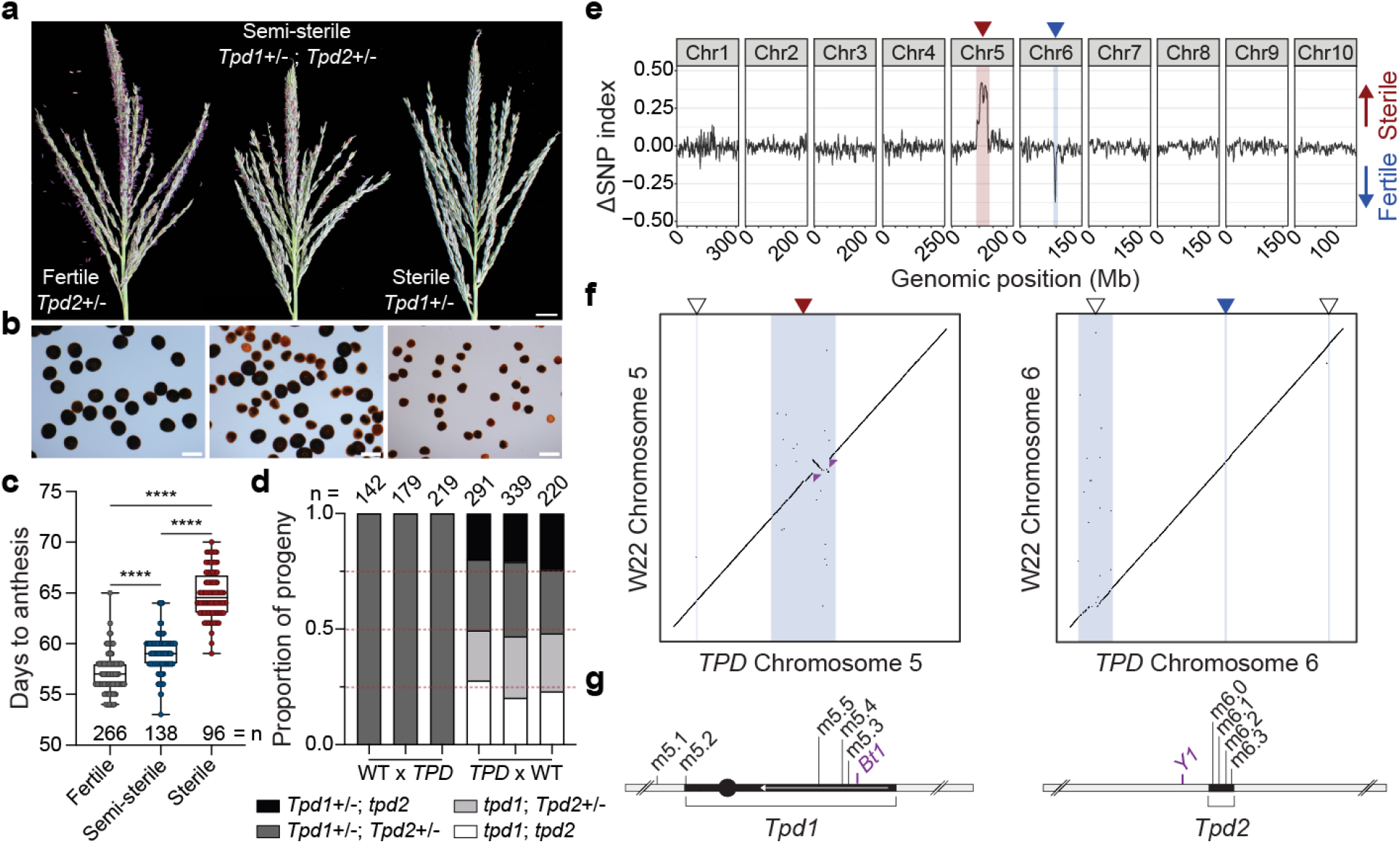
**A toxin-antidote system introduced from *mexicana* on chromosomes 5 and 6. a**, Representative tassels from fertile, semi-sterile, and sterile plants in a maternally segregating population. Examples shown are from synchronized planting dates. Scale bar = 1cm. **b**, I2KI viability staining of pollen from *Tpd2*+/-, *Tpd1*+/-; *Tpd2*+/-, and *Tpd1*+/- plants. Scale bar = 0.1mm. **c**, Measurement of days to anthesis in fertile, semi-sterile, and sterile phenotypic classes. Data are from three independent field populations. **** p < 0.0001 (Two-tailed Mann-Whitney test). **d**, Genotypic segregation ratios in reciprocal crosses. Genotypes were defined using codominant molecular markers. *n* represents the sample size for each progeny population. The red dashed lines denote a perfect 1:1:1:1 genotypic segregation ratio. Normal segregation is only observed in maternal progeny. **e**, Bulk segregant analysis (BSA) of fertile and sterile progeny pools indicates that *Tpd1* (red arrow) is necessary and sufficient for dominant male sterility (toxin) while *Tpd2* (blue arrow) is associated with fertility (antidote). FDR ≤ 0.01 (Benjamini-Hochberg method). **f**, Dot plots of chromosomes 5 and 6 showing multiple alignment between the *TPD* and W22 reference genomes. The blue lines and shaded regions correspond to five fully scaffolded intervals of *mexicana* introgression (indicated by arrows). As in (e), the red and blue arrows mark the *Tpd1* and *Tpd2* intervals, respectively. The small purple arrows indicate breakpoints of a ∼13Mb paracentric inversion present within the *Tpd1* haplotype on 5L. **g**, Summary graphics depicting the *Tpd1* and *Tpd2* intervals, as well as associated markers. The 13Mb inversion is indicated as a reverse arrow.

Genetic mapping placed *Tpd1* in a large interval surrounding the centromere of chromosome 5, while *Tpd2* was placed in a 1.5Mb interval on 6L (Extended Data Fig. 1c,d).

The selfish nature of *TPD* led us to liken it to previously described genic meiotic drivers that operate via post-meiotic gamete killing^27,29,31^. These systems generally encode a toxin (or distorter) that acts *in trans* to disrupt proper reproductive development. Only gametes containing a cell-autonomous antidote (or resistant responder allele) can suppress these effects in a gametophytic manner. While the toxin was clearly encoded by *Tpd1*, the *TPD* system was unusual in that it featured a genetically unlinked antidote, namely *Tpd2*. However, the absence of *tpd1; Tpd2* recombinants in the progeny of W22 x *TPD* crosses argued that *Tpd2* alone was insufficient for suppression of pollen abortion (Figure 2d; Supplementary Table 2). We reasoned that this might reflect the additional requirement for another antidote, linked to *Tpd1*, that could explain the observed rate of pollen abortion (∼75%). Linked modifiers in drive systems are common and generally ascribed to the co- evolutionary struggle between distorters and rapidly accumulating suppressors^14,29^.

SNP genotyping of the two homozygous lines identified 13 *mexicana* introgression intervals, 7 of which were shared between backcross generations (Extended Data Fig. 2a). As predicted from the single pollen sequencing data, the highest regions of SNP density were present on chromosomes 5 (*Tpd1*) and 6 (*Tpd2*), coinciding with *Bt1* and close to *Y1*, respectively (Extended Data Fig. 2a).

However, other regions strongly overrepresented in homozygous progeny were only partially overrepresented in *TPD* pollen, including additional peaks on 5S, 6S and 6L (Extended Data Fig. 2b). This likely reflected the presence of recombinant pollen grains that competed poorly during pollination.

To determine gene content in these and other introgression intervals, we performed *de novo* genome assembly from homozygous *Tpd1;Tpd2* BC8S3 seedlings using Oxford Nanopore long sequencing reads (∼30Kb read length N50). Repeat-graph based assembly^45^ was paired with long and short read polishing^46^ to yield a contig-level primary assembly. We then scaffolded the contigs with Dovetail Omni-C chromatin contact data to obtain a chromosome-level assembly (see Methods; Supplementary Table 3) with fully scaffolded *mexicana* introgression intervals on chromosomes 5 and 6 (Figure 2f). We noted the presence of a 1.9Mb *mexicana* introgression interval on 5S linked to the *Tpd1* haplotype and strongly over-represented in both our bulk sequencing and single-pollen grain data (Figure 1h,i; Figure 2f). Within this interval, we identified 10 genes with expression in pollen, one of which, *Dicer-like 2* (*Dcl2*), had excess nonsynonymous substitutions within conserved domains (Figure 3a), suggesting the possibility of adaptive evolutionary change^47^. Remarkably, absolute genetic linkage (n=214) between this locus (hereafter *dcl2^T^*) and *Tpd1* was conditioned on passage through the male germline from heterozygous *TPD* plants, while recombination between *dcl2^T^*and *Tpd1* occurred at the expected frequency (∼12%) when crossed as female (Figure 3b). This was very strong evidence for a linked antidote and likely explained the maintenance of this interval across thirteen backcross generations.

**Figure 3.**
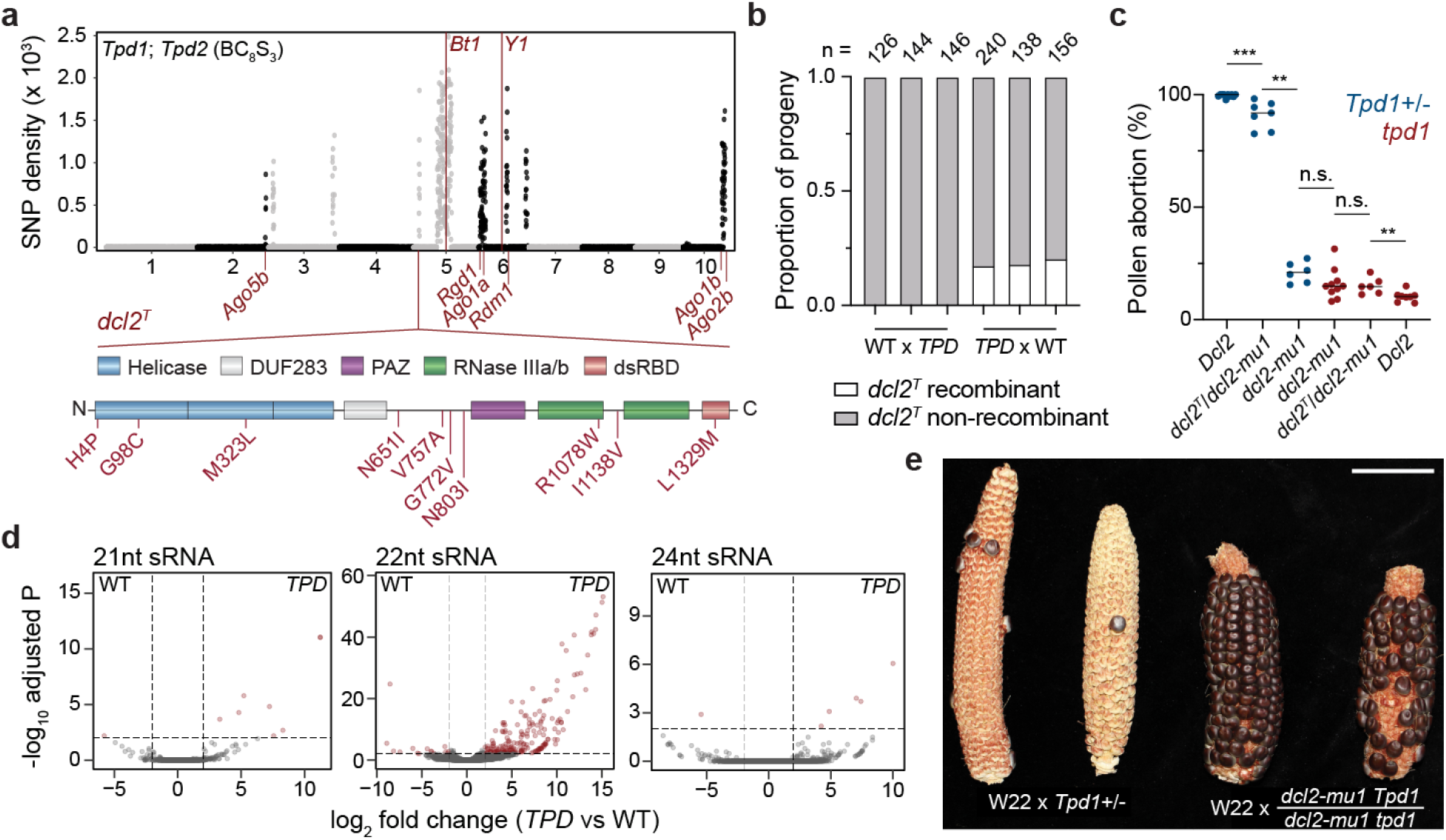
***Dicer-Like 2* from teosinte is a linked antidote for toxic 22nt small interfering RNA. a**, Genome-wide *mexicana* SNP density in bulk sequenced *Tpd1; Tpd2* (BC8S3) plants. A subset of *mexicana* introgression intervals (in addition to *Tpd1* and *Tpd2*) are selectively maintained and encode RNAi factors (*Dcl2*, *Ago1*, *Ago2*, *Ago5*, *Rgd1/Sgs3* and *Rdm1*). A *mexicana*-derived allele of *Dcl2* (*dcl2^T^*) with a high rate of nonsynonymous substitution is maintained in linkage to *Tpd1*. **b**, Rates of recombination between *dcl2^T^*and *Tpd1* in replicate reciprocal crosses. *dcl2^T^* exhibits tight pseudolinkage with *Tpd1* when propagated as male (0cM), but not as female (18.7 ± 1.6 cM). *n* represents the sample size for each progeny population. **c**, Measurements of pollen viability in *Tpd1/tpd1* and *tpd1* plants containing combinations *Dcl2*, *dcl2^T^*, and *dcl2-mu1*. Addition of the *dcl2- mu1* hypomorphic allele is sufficient for suppression of *Tpd1*-mediated pollen abortion, implicating 22nt siRNAs as the toxin underlying drive. Data points correspond to measurements from individual plants (*n =* 6-10). ** p < 0.01; *** p < 0.001 (Two-tailed Mann-Whitney test). **d**, Volcano plots showing 21nt (*n* = 9), 22nt (*n* = 212), and 24nt (*n* = 6) sRNA clusters that are differentially expressed in wild type and *TPD* pollen. The accumulation of ectopic 22nt siRNAs occurs in *TPD* pollen specifically. Log2 fold change ≥ 2, FDR ≤ 0.01. **e**, Representative ears from replicate crosses containing wild type *Dcl2* (W22 x *Tpd1/tpd1*) or *dcl2-mu1* (W22 x *dcl2-mu1 Tpd1/ dcl2-mu1* tpd1) in linkage to *Tpd1.* Pollen parents homozygous for *dcl2-mu1* restore seed set. Scale bar = 4cm.

*Dcl2* encodes a Dicer-like protein responsible for the production of 22nt small interfering RNAs (siRNAs) from hairpins, as well as secondary small RNAs from double-stranded RNA (dsRNA) templates produced by the coordinated action of RNA-DEPENDENT RNA POLYMERASE 6 (RDR6) and SUPPRESSOR OF GENE SILENCING 3 (SGS3)^48^. In *Arabidopsis thaliana*, DCL2 function is superseded by DCL4 and endogenous levels of 22nt siRNAs are low^49^. However, DCL2 can fulfill roles in silencing and antiviral immunity when DCL4 function is lost^49,50^, sometimes resulting in “toxic” pleiotropic defects associated with gene targets of 22nt siRNAs^49,51,52^. These observations stem from the unique biological properties of 22nt siRNAs, which are responsible for propagation of systemic silencing signals that move between cells^53^ and transitive amplification of silencing both *in cis* and *in trans*^54^. In *dcl2^T^,* non-synonymous changes were clustered within the DExD/H RNA Helicase domain of Dicer (Figure 3a), which has been shown to alter substrate preference and processing efficiency of dsRNA, but not hairpin RNA, in both plants and invertebrates^55–57^.

To explore the role of 22nt siRNAs in the *TPD* phenotype, we tested mutants in 22nt siRNA biogenesis for their ability to act as antidotes. We isolated maternal *dcl2^T^* recombinants and compared them to the *dcl2-mu1* allele in the W22 inbred background, which has a *Mu* transposon insertion in the 5’ untranslated region (5’ UTR), 200bp upstream of the start codon. In *dcl2^T^*/*dcl2-mu1 Tpd1*, pollen abortion was partially suppressed, while pollen from *dcl2-mu1/dcl2-mu1 Tpd1* plants were almost fully viable (Figure 3c). This meant that stacking over the *dcl2^T^* allele had a synergistic effect, strongly supporting its role as a partial antidote, and indicating that the sporophytic production of 22nt siRNAs in diploid meiotic cells was responsible for the *TPD* phenotype. To test the idea that 22nt siRNAs might be responsible for *TPD*, we sequenced pollen small RNAs from *TPD* and WT siblings, and found that while sRNA composition was similar overall, the *Tpd1* haplotype triggered a strong, 22nt-specific response (Figure 3d). Consistent with these 22nt small RNAs being responsible for the *TPD* phenotype, we observed almost complete rescue of sterility in *dcl2-mu1/dcl2-mu1 Tpd1/+* pollen parents (Figure3e). Intriguingly, several other introgression intervals observed in one or the other backcross individual also included genes encoding components of the small RNA biogenesis pathway, including *ago1a*, *ago1b,* and *rgd1,* the homologue of SGS3 (Figure 3a; Extended data Fig. 2a). These intervals were also observed in single pollen grain sequencing along with *dcl2^T^* (Extended Data Fig. 2b). To determine if these genes were also capable of acting as an antidote, we crossed mutants in *rgd1* to *TPD* plants. Segregation of *rgd1* in the germline of heterozygotes resulted in close to 50% viable pollen (Extended Data Fig. 2c), suggesting that it functions as a cell autonomous gametophytic suppressor in a manner similar to *Tpd2*. We concluded that mutants in primary 22nt small RNA synthesis (*dcl2-mu1*) blocked production of the toxin, while mutants in secondary 22nt small RNA synthesis (*dcl2^T^* and *rgd1*), and potentially in small RNA function (*ago1a,b*), acted as antidotes.

### Germline small RNA target the pollen lipase gene *Teosinte Drive Responder*

In order to identify the origin, and the targets of Dcl2-dependent small RNAs, we performed small RNA sequencing from WT, *dcl2^T^*, and *dcl2-mu1* plants. Analysis revealed that 22nt siRNAs were the dominant species in WT pollen (Extended Data Fig. 3a,b) and defined 804 high-confidence 22nt siRNA pollen-specific clusters (log2 FC ≥ 2, FDR ≤ 0.01) (Supplementary Table 8). . As expected, these clusters depended on *Dcl2* (p < 0.0001, ANOVA) and there were even fewer 22nt siRNAs in *dcl2-mu1* than in *dcl2^T^* (Extended Data Fig. 3c). Remarkably, over half (54.6%) of all pollen-specific 22nt species were derived from endogenous hairpin precursors (hpRNAs) (Extended Data Fig. 3d,e,g). Hairpin short interfering RNAs (hp-siRNAs) were disproportionately 22nt-long, derived from a single strand (Extended Data Fig. 4a,b) with high thermodynamic stability (Extended Data Fig. 4c,d). Based on these criteria (and a minimum expression cut-off), we identified 28 hp-siRNA producing loci in the genome, with at least one hairpin on every chromosome except chromosome 4 (average 2.1 ± 1.3 per chromosome). hpRNAs can serve as a powerful means to silence transposons^58^, and 22nt siRNAs targeting *Gypsy* and *Copia* LTR retrotransposons were abundant in pollen, as were those targeting *Mutator* and *CACTA* elements (Extended Data Fig. 3d). Surprisingly, we also found evidence for pollen-specific silencing of at least 30 protein coding genes (Extended Data Fig. 3d,f,g). Germline-specificity is a common feature in *SD* systems, as such factors can avoid the evolutionary conflicts imposed by pleiotropic fitness defects in the diploid stage of the life cycle^34^.

In *TPD* pollen, we observed the accumulation of 158 ectopic 22nt siRNA clusters across the genome (log2 FC ≥ 2, FDR ≤ 0.01) (Supplementary Table 9), and a general up-regulation of genes associated with 22nt siRNA biogenesis and function (Extended Data Fig. 5a). Nearly 60% of all ectopic 22nt siRNAs in *TPD* pollen targeted TEs of the *P Instability Factor* (PIF)/Harbinger superfamily (Extended Data Fig. 5b), whose expression was *TPD* specific (Extended Data Fig. 5c-e). Interestingly, this superfamily is also activated following intraspecific hybridization and anther culture in rice^59^. However, a subset of protein-coding genes was also targeted in *TPD* pollen specifically (Extended Data Fig. 5b). Given that a reduction in 22nt siRNAs suppressed the *TPD* phenotype, we hypothesized that inappropriate silencing of these genes might disrupt male reproductive development. In total, we identified 4 genes that gained ectopic 22nt siRNAs in *TPD* pollen, ∼62% of which came from a single gene (Zm00004b012122) also located on chromosome 5S (Extended Data Fig. 6a). Relative to other targets, this gene exhibited highly specific expression in pollen (Extended Data Fig. 6b,c). Zm00004b012122 encodes a GDSL triacylglycerol (TAG) lipase/esterase, defined by a core catalytic sequence motif (GDSxxDxG), with roles in lipid metabolism, host immunity, and reproductive development^60^. In maize, both *male sterile 30* (*ms30*) and *irregular pollen exine 1* (*ipe1*) mutants disrupt genes encoding a GDSL lipase and are completely male sterile^61,62^. Similar functions have also been reported in rice^63^ and *Arabidopsis*^64^, where germline-specific GDSL lipase activity is essential for proper aliphatic metabolism in developing anthers, and loss of function results in aberrant pollen exine development.

DCL2-dependent 22nt siRNAs engage primarily in translational repression of their targets^65^, and consistently all 4 target genes had similar or higher levels of mRNA in *TPD* pollen (Extended Data Fig. 6c). We raised antiserum against the GDSL lipase for immunoblotting, choosing a surface exposed peptide located between putative pro-peptide processing sites reflecting endoplasmic reticulum (ER) localization^62^. The GDSL lipase protein accumulated strongly in both 5mm anthers and mature pollen from WT plants, but was absent from leaf and from *TPD* anthers and pollen, supporting the conclusion that 22nt siRNAs mediate translational repression (Extended Data Fig. 6d). Further, whole protein extracts from *TPD* anthers had reduced esterase activity which was ameliorated in pollen containing *Tpd2* but not in pollen with *Tpd1* alone (Extended Data Fig. 6e).

Gene ontology analysis of genes upregulated in WT and *TPD* pollen strongly supported translational suppression of the GDSL lipase as the primary cause of developmental arrest and abortion of pollen in *TPD* plants (Extended Data Fig. 6f,g). Finally, mRNA expression began post-meiotically at the 3mm (tetrad) stage, peaking in 5mm anthers and mature pollen (Extended Data Fig. 7a). This expression pattern was conspicuously similar to the developmental window in which *TPD* pollen abortion begins (Figure 1f), suggesting that this gene might act as a ‘responder’ to *Tpd1-*driven distortion. Based on all these observations, we defined Zm00004b012122 as the primary candidate for targeting by *Tpd1* toxin activity, renaming it *Teosinte Drive Responder 1* (*Tdr1*).

### Teosinte-Specific Hairpin small RNAs Target *Tdr1* and trigger pollen abortion

As ectopic silencing at protein-coding genes only occurred in the presence of the *Tpd1* haplotype, we reasoned that the distorter must generate small RNAs capable of triggering silencing *in trans*. In plants, microRNAs (miRNAs), secondary siRNAs, and hp-siRNAs all have this capacity^66^. Processed small RNA duplexes are loaded into ARGONAUTE (AGO) proteins, passenger strands are released, and RNase H-like slicing activity is targeted by guide strand homology, as is translational repression^67^. Silencing can be amplified via the coordinated action of RDR6 and SGS3^54^. RNase H-mediated slicing results in an exposed 5’ phosphate that allows for ligation of 3’ cleavage products^68^. Using an improved degradome sequencing technique in *TPD* pollen, iPARE-seq (Roche et al., in prep), we could identify putative cleavage sites responsible for triggering silencing at the *Tdr1* locus (Figure 4a,b). We simultaneously searched for non-coding RNA within the *Tpd1* haplotype that produced 22nt sRNAs capable of triggering silencing. This approach yielded only one candidate; a large hpRNA similar to those identified previously in WT pollen (Figure 4c). Interestingly, this hairpin was uninterrupted in the *mexicana*-derived *Tpd1* interval, and produced high levels of *TPD*- specific 22nt hp-siRNAs (Figure 4d,e). In the W22 genome, we identified two large transposon insertions that interrupted this locus, which produced no small RNA, indicating that it was non-functional in maize, consistent with being responsible for *TPD* (Figure 4c). By comparison with centromere placement in other maize inbreds^69^, the hairpin is on the short arm of chromosome 5, 5Mbp from the centromere.

**Figure 4.**
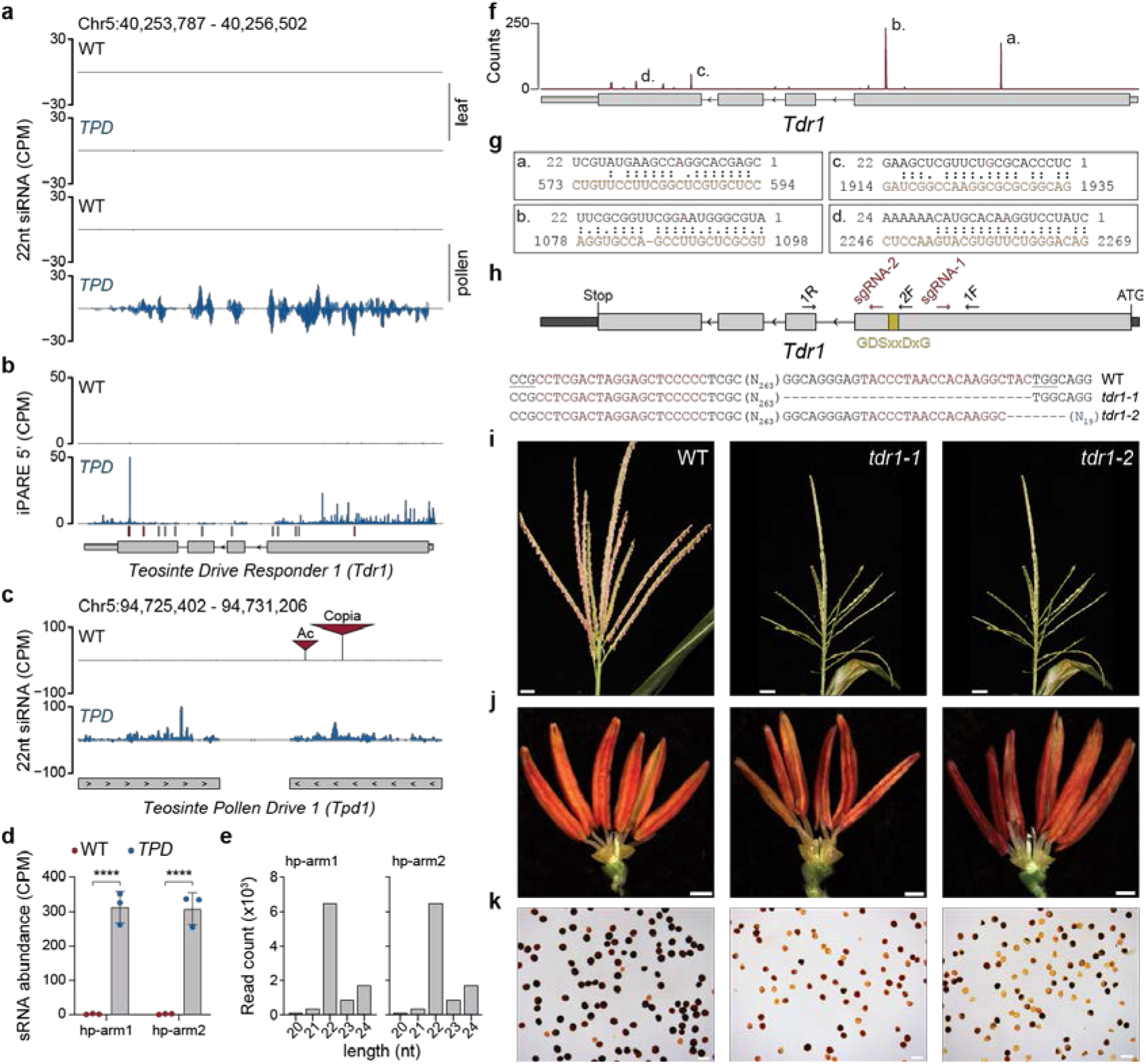
**22nt siRNAs from a *mexicana*-derived hairpin (*Tpd1*) target *Teosinte drive responder 1 (Tdr1),* an essential pollen gene. a**, Browser shots showing 22nt small interfering RNA (siRNA) levels at the *Tdr1* locus in leaf and pollen tissue from wild type and *TPD* genotypes. Ectopic 22nt siRNAs accumulate in *TPD* pollen specifically. **b**, iPARE-seq depicting the accumulation of 3’-OH cleavage products at the *Tdr1* locus. The 5’ nucleotide of each fragment is plotted and corresponds to the position of Argonaute mediated cleavage. Tick marks indicate predicted target sites for hairpin siRNAs (hp-siRNAs) derived from the *Tpd1* hairpin. Sites with (red) and without (gray) iPARE read support are shown. **c**, Browser shot of 22nt hp-siRNA accumulation at the *Tpd1* hairpin. The hairpin locus is disrupted by transposable element insertions in the W22 genome. Data shown (**a**, **b**, **c**) are normalized counts per million (CPM). **d.** 22nt hp-siRNA abundance at the *Tpd1* hairpin locus in WT and *TPD* pollen. n = 3 replicates per condition. **** p < 0.0001 (Mann-Whitney test). **e** Average size distribution of reads mapping to the *Tpd1* hairpin. **f**, small RNA target site prediction at the *Tdr1* locus using psRNATarget. Counts indicate unique hp-siRNAs from *Tpd1* that target each site. **g,** Homology between the guide strand (black) and target strand (orange) is shown for the four most abundant hp-siRNAs. The 10^th^ (red) and 11^th^ nucleotides in the guide strand flank the site of Ago-mediated cleavage. *Tpd1 hp-siRNA b* is predicted to suppress translation. **h**, Schematic showing CRISPR/Cas9 targeting of the *Tdr1* locus. Edits corresponding to *tdr1-1* and *tdr1-2* alleles (blue) are shown. **i**, Developmentally synchronized tassels from wild-type and *tdr1* mutant T0 plants. *tdr1* mutants exhibit severely delayed anthesis. Scale bar = 3cm. **j**, Mature 5mm anthers from wild-type and *tdr1* mutant T0 plants. Scale bar = 1mm. **k**, I2KI viability staining of pollen from wild-type and *tdr1* mutant T0 plants. Scale bar = 0.1mm.

Target site prediction using psRNATarget^70^ uncovered 4 abundant hp-siRNA species predicted to target the *Tdr1* transcript *in trans* (Figure 4f). Three of these began with 5’-C indicating loading into Ago5, and had iPAREseq support indicating cleavage of *Tdr1* (Figure 4g). However, the most abundant hp-siRNA, *Tpd1-*siRNAb, was 22nt in length and began with 5’-A indicating loading into Ago2 (Figure 4g). *Tpd1-*siRNAb has an asymmetric bulge predicted to enhance silencing transitivity and systemic spread between cells^71,72^, and had only limited iPAREseq support indicating translational repression (Figure 4b). In order to confirm that silencing of *Tdr1* was responsible for the *TPD* phenotype, we generated two independent frameshift alleles within the catalytic domain using CRISPR-Cas9 (Figure 4h). Homozygotes for *tdr1-1* and *tdr1-2* had identical male sterile phenotypes, with extensive pollen abortion that phenocopied *Tpd1* (Figure 4i-k).

Expression of the *Tpd1* hairpin was observed pre-meiotically in 1-3mm anthers, as well as in microspores (4mm anthers) where expression of *Tdr1* was first detected, but not in mature pollen (Extended Data Fig. 7b,c). According to published single cell RNA-seq data from developing maize pollen^73^, *Dcl2* is also expressed pre-meiotically consistent with its role in generating hp-siRNA from *Tpd1* (Extended data Fig. 7d). *Dcl2* is not expressed in microspores, but is expressed in mature pollen consistent with an additional function in production of secondary small RNAs from *Tdr1* (Extended data Fig. 7d). These results indicate a sequential order of events, in which expression of *Tpd1* pre- meiotically deposits small RNAs in microspores where they target *Tdr1*. Subsequent expression of *Dcl2* in mature pollen then promotes secondary small RNA production and translational suppression. Identification of *Teosinte Drive Responder 1* provided insight into the function of *Tpd2*. *Tpd1* hp-siRNAs were unaffected by *Tpd2*, which was instead required to suppress secondary small RNA biogenesis from *Tdr1,* along with the *mexicana* allele of *Dcl2,* namely *dcl2^T^* (Extended Data Fig. 8a). This indicates that *Tpd2* and *dcl2^T^* have additive effects on suppressing secondary small RNAs, consistent with their role as partial antidotes. While the molecular identity of *Tpd2* remains unknown, the 1.5Mb *Tpd2* interval contains 6 pollen-expressed genes in W22 (Extended Data Fig. 8b).

Intriguingly, one of these genes encodes the maize homologue of *Arabidopsis* RNA-DIRECTED DNA METHYLATION (RDM1), a critical component of the RNA-directed DNA methylation (RdDM) pathway^74^. This gene is significantly overexpressed in *TPD* pollen (Extended Data Fig. 8b), and it is possible that increased activity of RdDM might compete with the production of secondary small RNAs^75,76^, though further experimentation is required to support this conclusion (Extended Data Fig. 9).

### *Teosinte Pollen Drive* and RNA interference contributed to maize evolution and domestication

Population-level studies of both *mexicana* introgression lines and maize landraces have consistently identified an uninterrupted *mexicana*-derived haplotype surrounding the centromere of chromosome 5^41,77^. In landraces, this haplotype consistently showed high rates of linkage disequilibrium (LD), suggesting a virtual absence of recombination^77^. Consistent with reduced recombination, fine-mapping of *Tpd1* yielded very few informative recombinants (21/7,549) and none at all proximal to the hairpin (Extended Data Fig.1c). Comparative analysis of the *TPD* and W22 genomes revealed three megabase-scale inversions, one of which corresponded to a 13Mb event within the *Tpd1* haplotype and including *Bt1* on chromosome 5L (Figure 2f,g). The presence of this inversion, along with its physical proximity to the centromere, explained our mapping data (Extended Data Fig. 1c), and strongly suggested that the *Tpd1* haplotype behaves as a single genetic unit. Large structural variants, such as inversions, can reinforce genetic barriers between populations by contributing to the fixation of deleterious or incompatible variants^78^, and are frequently co-opted during drive evolution to ensure the integrity of distorter-responder pairs, and of linked bystander genes not involved in drive^14^. Functional analysis of variation in the *Tpd1* interval revealed a handful of putative loss-of-function mutations (n = 12) owing primarily to frameshifts, as well as high rates of nonsynonymous substitution (n = 277) . This result was in line with the estimates of increased genetic load within regions of low recombination^79^, and mirrored observations in other drive systems, which often accumulate deleterious recessive mutations in linkage^14^. In addition to coding sequence mutations, we found extensive structural variation in the form of 1519 high confidence insertions and deletions (indels) of various sizes (mean size = 5,419bp).

Intriguingly, the 13Mb paracentric inversion in the *Tpd1* haplotype (W22 Chr5:115,316,812- 124,884,039) almost entirely encompasses so-called “region D” adjacent to centromere 5 (W22 Chr5:118,213,716-126,309,970), that has undergone a dramatic domestication sweep in all maize inbreds relative to teosinte^69^. This region includes *Bt1*, which undergoes drive in the TPD system (Extended Data Fig. 1c). Hybridization between teosinte *mexicana* and early cultivated maize was a critical step in the adaptation and geographical dispersal of modern maize, following initial domestication from teosinte *parviglumis* in tropical lowlands^41,80,81^. We rationalized that, if *TPD* exists as a cryptic drive system in *mexicana*, signatures of drive might be found in traditional Mexican varieties (often called “landraces”^41^), and in sympatric *mexicana* populations in the central highlands of Mexico where hybridization still occurs. Our synthetic hybrids with maize inbred W22 retained approximately 13 intervals of the *mexicana* genome that persisted in serial backcrosses (Extended Data Fig. 2a,b), and we looked for signatures of these intervals in 256 accessions from 10 different traditional varieties of maize, along with their sympatric populations of wild *mexicana*^41^. We determined the mean frequency of *mexicana*-like alleles from each interval in each population from existing genotyping data^81^ and determined the Spearman’s rank correlation coefficient for each pair of intervals. We found that, in *mexicana* populations, the intervals encoding *Tpd1* (chr 5.79), *Tpd2* (chr 6.98) and *Rgd1* (chr 6.3) were highly correlated (Figure 5a), suggesting that this combination of toxin and antidote loci is ancestral in these *mexicana* populations. In traditional maize varieties, on the other hand, significant correlations were observed between 11 of the 13 intervals (Figure 5a). 3 of these intervals are tightly linked to genes encoding Argonaute proteins, specifically *Ago1a*, *Ago1b* and *Ago2b*, all of which are expressed in the male germline (Extended Data Fig. 2). A fourth Argonaute gene, the anther-specific *Ago5b*, is tightly linked to another of the 13 intervals retained in backcross hybrids (Extended Data Fig. 2a), but not in extant traditional maize varieties (Figure 5a). According to 5’ nucleotide analysis these Argonaute proteins are predicted to bind *Tpd1*-hp-siRNAa- d (Ago2 and Ago5), as well as secondary *Tdr1* 22nt siRNAs (Ago1), and it is conceivable that hypomorphic alleles could also act as partial antidotes in combination with *Tpd2*. In addition to intervals encoding *Dcl2*, *Rdm1* and *Rgd1/Sgs3*, this means that 7 of the 13 intervals are tightly linked to genes required for RNAi. These correlations suggest there has been strong selection on all of these modifiers to ameliorate the toxic effects of *Tpd1*, resulting in apparent gene drive.

**Figure 5.**
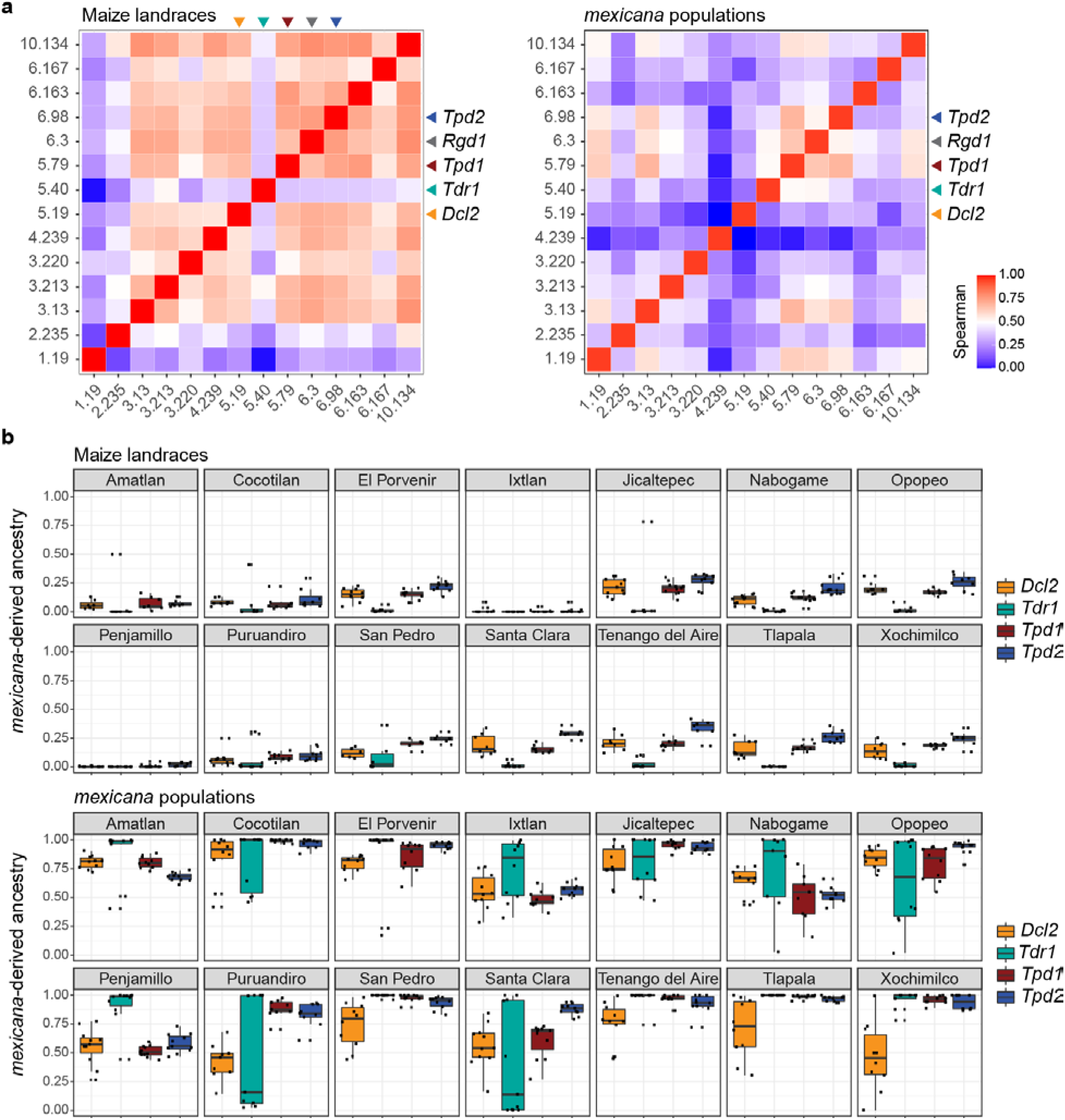
Signatures of *Teosinte Pollen Drive* in modern maize, maize landraces and sympatric *mexicana*. a, Frequency of *mexicana*-derived alleles were calculated for 1 Mb intervals associated with *TPD* on chromosomes 1,2,3,4,5,6 and 10. Correlations are shown between population means from each of 14 maize landraces (left) and sympatric *mexicana* populations (right). Intervals on chromosomes 5, 6 and 10 include *Dcl2* (5.19), *Tdr1* (5.40), *Tpd1* (5.79), *Rgd1/Sgs3* (6.3) and candidate genes *Ago1a* (6.3), *Tpd2/Rdm1* (6.98), *Ago1b* and *Ago2b* (10.134). Correlations were observed for most of the intervals in maize landraces, except for *Tdr1 (green arrow),* but only for intervals including *Tpd1, Rgd1* and *Tpd2* in *mexicana*. Spearman correlation coefficients are displayed as a heatmap. **b**, *mexicana*-derived ancestry in each of 14 maize landraces (above) and sympatric *mexicana* populations (below) in *Dcl2*, *Tdr1*, *Tpd1* and *Tpd2* intervals. The *Tdr1* interval (green) is monomorphic in most of the maize landraces, but shows extreme dimorphism in 7 out of 14 sympatric *mexicana* populations.

In contrast to RNAi genes, variation at *Tdr1* displays no such correlation with the co-inherited intervals in traditional maize varieties (Figure 5a). In fact, *Tdr1* is strongly monomorphic in traditional maize varieties, while in *mexicana Tdr1* displays extreme polymorphism (Figure 5b). We considered the possibility that this locus has evolved to become immune to silencing in modern maize, a predicted outcome of selfish genetic systems^14^. Remarkably, a recent survey of maize and teosinte genome sequences^82^ reveals that 3 of the 4 *Tpd1*-hp-siRNA target sites in *Tdr1* exhibit extensive polymorphism in maize and teosinte, including an in-frame deletion of the target site seed region for *Tpd1*-hp-siRNAa and a SNP at position 11 in target sites for *Tpd1-*hp-siRNAb, that are predicted to reduce or abolish cleavage and translational inhibition (Figure 6a). *TPD* pollinations of the temperate inbred B73, which carries the deletion haplotype, resulted in 50% sterile (44/83) and fully fertile (35/83) offspring in advanced backcrosses, as well as rare more highly sterile presumptive recombinants (4/83), consistent with these expectations. Surveys of the frequency of the deletion haplotype across *Zea* found it widespread, suggesting an origin prior to speciation of *Zea mays* from *Z. luxurians* and *Z. diploperennis* (Figure 6b), while it is absent from Z. *nicaraguagensis* and *Tripsacum dactyloides*. The frequency of the deletion haplotype is relatively low in *mexicana* (12%) compared to *parviglumis* (46%), and increases in tropical maize, traditional maize varieties (landraces), popcorn and inbreds, where it is nearly fixed in several modern inbred groups (98%), suggesting a trajectory of spread to North and South America.

**Figure 6.**
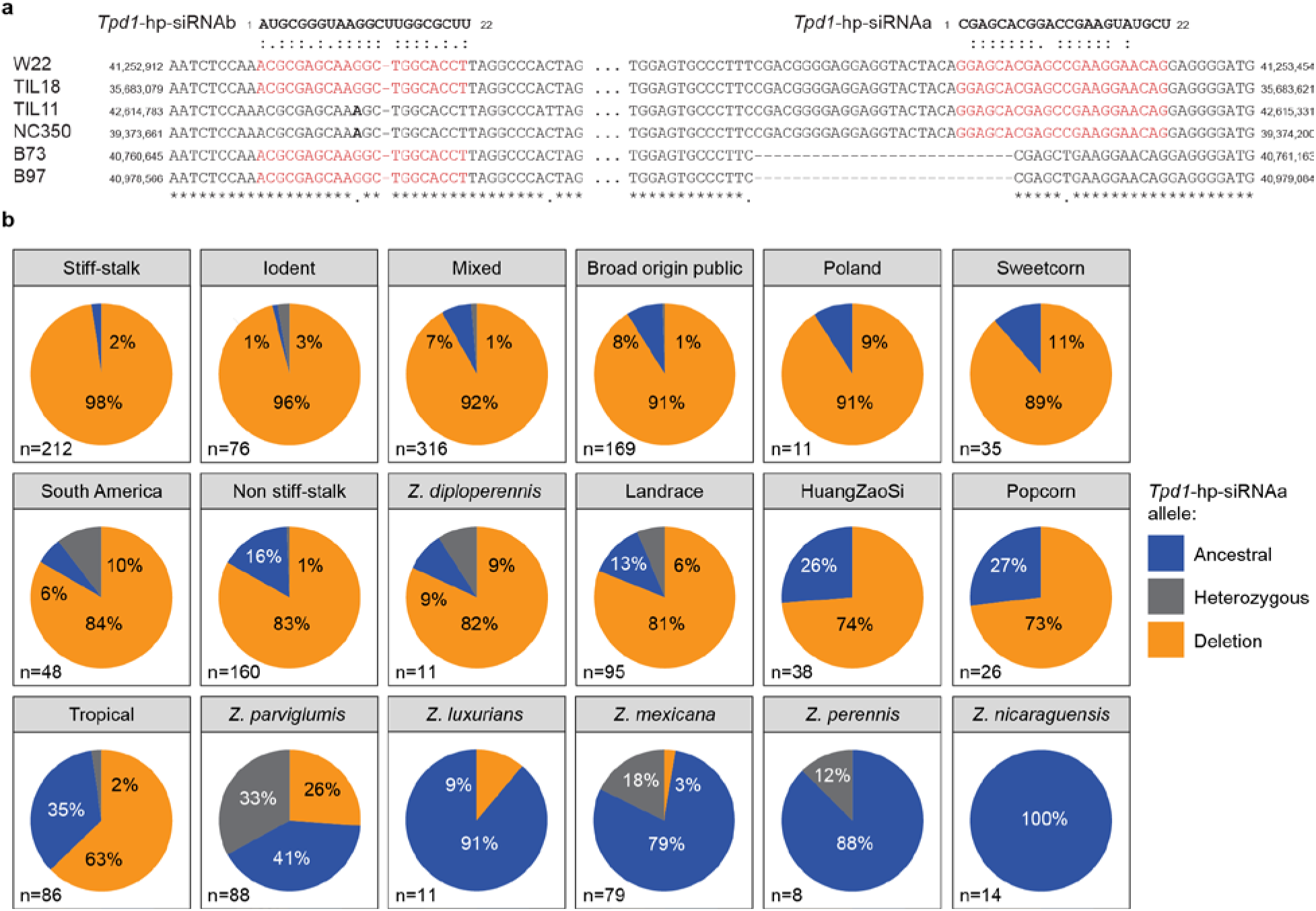
***Tpd1* hp-siRNA target site deletion in *tdr1* has spread to modern maize from teosinte. a**, Sequence complement of the *Tpd1-hp*-siRNA a and b target sites in *Tdr1*, indicating a 27bp in-frame deletion found in modern maize, maize landraces and in teosinte, that removes the *Tpd1*-hp-siRNAa seed sequence, and a SNP on the 11^th^ nucleotide of *Tpd1*-hp-siRNAb that is predicted to reduce binding. **b**, Pie charts indicating frequency of the deletion in 1,483 resequenced genomes from maize and teosinte ^85^, aligned with the B73 reference genome (GATK 3.0). The deletion allele (blue) arose in teosinte and quickly spread through maize landraces in Central and South America, before fixation in modern stiff stalk, but not tropical maize inbred lines. High frequencies of heterozygosity in *mexicana* and *parviglumis* are consistent with recent or ongoing pollen drive.

To further characterize the evolution of genomic elements associated with *TPD*, we applied the haplotype-based statistic iHS^83^ and the FST statistic^84^ to identify genome-wide signals of natural selection (see Methods) using variant data for *parviglumis*, *mexicana*, landraces, and modern lines^85,86^. We considered values in the top 5% of the genome-wide empirical distributions of these statistics as significant. FST was effective at identifying known domestication loci, but did not show significant signals of selection for the *TPD*-linked genes (Supplementary Table 4). However, we found that *dcl2*, *tdr1*, *rdm1* and the hairpin region contained significant SNPs in teosinte based on iHS (Supplementary Table 5), indicating these loci may be under selection. More conservative analyses using Bonferroni *p*-value adjustment for the multiple tested populations and windowed iHS statistics found signatures of selection in *dcl2* for *mexicana* and for the hairpin region in *parviglumis* (Supplementary Table 5).

## Discussion

### *Teosinte Pollen Drive* (*TPD*) is a selfish genetic system that defies Mendelian inheritance

Like other selfish systems, *TPD* contributes to hybrid incompatibility, in this case between maize and teosinte. Unlike teosinte crossing barriers *tcb-1*, *Ga-1* and *Ga-2*^87^, which prevent fertilization, *TPD* resembles Bateson-Dobzhansky-Muller (BDM) incompatibility (also known as DMI) in that it acts post zygotically, resulting in sterile progeny. In canonical BDM, however, hybrid lethality is due to the unmasking of duplicate deleterious alleles, so that backcrosses to either parent are fully fertile. In *TPD*, backcrosses to maize, but not to teosinte, result in pollen abortion no matter how many backcross generations are observed. This is because *TPD* is a special case of BDM that operates via a toxin-antidote model and results in meiotic drive. It has been proposed that in order for gamete killers to achieve drive at the population level, they must compensate somehow for loss of fecundity^88^. In principle, however, maize-teosinte hybrids are extremely vigorous with numerous tassels, so that these wind-pollinated species may be less sensitive to reductions in male fertility. This is especially true during domestication, when early domesticates are typically less prolific than wild relatives, and at lower population size. In such circumstances, gene drive from wild relatives into domesticates could have an outsized impact on hybridization. Furthermore, the presence of linked regions of potential selective advantage, such as the inversion spanning “region D”, or alleles involved in flowering time, might provide some compensation during domestication.

*Tpd1* encodes a long non-coding hairpin RNA that produces 22nt hp-siRNAs that accumulate in the male germline. These hp-siRNA kill subsequent pollen grains by targeting the genetically linked responder gene *Tdr1* (Extended Data Fig. 9a, b). This effect is countered by at least two gametophytic antidotes, a linked hypomorphic allele of *Dcl2* and the unlinked *Tpd2* locus on chromosome 6 (Extended Data Fig. 9c). The genetic architecture of this system, consisting of multiple linked and unlinked loci, deviates from previously established toxin-antidote systems. In rice, for instance, the *qHMS7* quantitative trait locus (QTL) is a selfish genetic element composed of two tightly linked open reading frames (ORFs)^89^. Similarly, the *wtf4* driver in *S. pombe* features two alternatively spliced transcripts derived from the same locus^31^. In contrast, the *Tpd1* haplotype results in tight pseudo-linkage between *Tpd1*, *Tdr1* and *dcl2^T^* but only when transmitted through the male.

While recombinants occur in single-pollen grains, they are not transmitted to the next generation (Figure 1), and maternal recombinants between *Dcl2^T^* and *Tpd1* are completely male sterile (Figure 2). These recombinants produce far more secondary 22nt small RNAs at *Tdr1* (Extended Data Fig. 8a), providing an explanation for the failure to transmit recombinants through pollen. *Tpd2* is unlinked but acts cell autonomously, so that independent assortment of *Tpd1* and *Tpd2* occurs in female gametes, but never in male, implying that gametophytic suppression of pollen killing requires co-segregation of *Tpd2* with *Tpd1*. A similar scenario was reported for drive in fission yeast mediated by *wtf13*. The emergence of a spontaneous unlinked suppressor, *wtf18-2,* was found to selectively suppress spore killing only in spores in which it is co-inherited with the toxin^90^. In both cases, the selective suppression of drive can be interpreted as selfish behavior on the part of the antidote. Such interactions arise repeatedly as part of the co-evolutionary struggle between drivers and the host genome, directly influencing patterns of co-segregation even in the absence of physical linkage.

Ultimately, this cycle of suppression and counter-suppression can be expected to result in complex, polygenic drivers that exist in a continuum of cryptic states (Extended Data Fig. 10).

The conspicuous maintenance of *mexicana* introgression intervals containing RNAi factors across many backcross generations supports this idea (Extended Data Fig. 2a; Extended Data Fig. 10). *mexicana* introgression was a key step in facilitating adaptation of early maize domesticates to the highland conditions of central Mexico^41,91^. In theory, the presence of a driver in *mexicana* could directly impact introgression by skewing patterns of heritability in hybrid populations. Genome scans of sympatric maize and *mexicana* have identified multiple regions of introgression associated with adaptive variation, some of which overlap with the genomic interval corresponding to the *Tpd1* haplotype^41^ and other intervals undergoing drive^92^, and we find that intervals associated with drive in pollen are significantly correlated with each other in maize landraces, but not in sympatric *mexicana* populations (Figure 5). We postulated that the most powerful suppressor of all would be an “immune” target gene, in which hp-siRNA target sites in *Tdr1* had been mutated. Remarkably, such in-frame “immune” haplotypes were found in wild taxa in *Zea*, and have been progressively fixed from tropical to temperate stiff-stalk maize inbreds (Figure 6), suggesting that *Teosinte Pollen Drive* may be an ancient system that has impacted admixture throughout the history of the genus, reaching fixation in modern maize. *Teosinte Pollen Drive* complements the hypothesized role of *Abnormal-10*, a chromosomal driver of female meiosis, that simulations suggest may have been responsible for the re-distribution of heterochromatic knobs in maize, *parviglumis* and *mexicana*^20,93^, potentially along with thousands of linked genes^21^.

Meiotic recombination has been proposed to be an evolutionary consequence of defense against drive, as the separation of individual drive components is an efficient way to purge selfish genetic elements^94^ and rare recombinants in the *SD* and *t*-complexes in mouse and Drosophila exhibit either impaired SD or suicidal male sterility^24,28^. As such, many drive complexes evolve in regions of reduced recombination and select for further structural rearrangements, some of which can give rise to patterns of variation that influence morphological traits not associated with drive itself^14^. The *t*- complex, which contains genetic loci governing SD, embryonic lethality, and morphological phenotypes such as tail length is a clear example^24^. At the structural level, the *Tpd1* haplotype is consistent with this notion, encompassing both the core centromeric region and a 13Mb paracentric inversion on chromosome 5L. In *mexicana*, the *Tpd1* haplotype captures upwards of 243 protein-coding genes including at least 12 putative loss of function alleles, significant levels of nonsynonymous substitution, and numerous small SVs. Independent studies have identified major QTLs within the centromeric region of chromosome 5 associated with flowering time^77^, hybrid yield^95^, and up to 6 domestication traits in teosinte-maize backcross populations^96^. This region also overlaps with introgression intervals associated with adaptive variation in traditional maize varieties^41,92^. Finally, the 13Mb inversion within the *Tpd1* haplotype encompasses one of largest known domestication sweeps between maize inbreds and teosinte^69^, accounting perhaps for the unusual size and rapid emergence of “Region D”. Signals of selection at *TPD* loci themselves, by contrast, were weaker than at known domestication loci, consistent with the idea that drive itself is not adaptive.

We show that DCL2-dependent 22nt small RNAs stemming from long hpRNAs can function as selfish genetic elements in pollen. In *Arabidopsis,* 22nt siRNA biogenesis is carefully regulated due to ectopic silencing of host genes^49,52,54,65^. Furthermore, 21-22nt siRNAs from Arabidopsis pollen target imprinted genes as well as transposons, and mediate triploid seed abortion^97,98^. In this context, the fact that DCL2 activity in the maize germline is unconstrained, producing highly abundant 22nt siRNAs that target mobile elements and protein-coding genes alike, is striking. Our observations are highly reminiscent of the testis-specific hairpin-RNA (hpRNA) pathway in *Drosophila melanogaster*^99,100^. In this system, silencing of protein-coding genes by a set of recently evolved hairpins is important for proper regulation of gene expression during male reproductive development^100^. In mammals, endo-siRNAs in the oocyte are generated from hairpin and antisense precursors by an oocyte specific Dicer isoform (*Dcr-O*) lacking the helicase domain, and have recently been found to have an essential function in global translational suppression^101–103^.

Importantly, the Winters sex-ratio (SR) distortion system in *Drosophila simulans* is suppressed by two hpRNAs, *Not much yang* (*Nmy*) and *Too much yin* (*Tmy*), which are essential for male fertility and sex balance^104,105^, while the “Paris” SR system may require a component of the piRNA pathway^106^. The remarkable parallels between all of these systems, and between Dcr-O and *Dcl2^T^*, which both have potential defects in the helicase domain, invites speculation that selection for selfish behavior is an efficient means by which germline small RNAs can propagate within a population.

Such propagation provides a plausible origin for “self”-targeting small RNAs in the germlines of plants and animals.

## MATERIALS AND METHODS

### Plant material and growth conditions

The *TPD* lineage traces to teosinte *mexicana* collected near Copándaro, Michoacán, Mexico in December 1993. Gamete a, plant 4 of collection 107 was used in an initial outcross to the Mid- western US dent inbred W22 and subsequently backcrossed. *Tpd1;Tpd2* (BC8S3) homozygous lines were used for whole-genome sequencing and *de novo* genome assembly. All additional experiments were performed using *Tpd1/tpd1; Tpd2/tpd2* (BC11-BC13) plants or populations derived from maternal segregation of these lines. The *lbl1-rgd1* and *dcl2-mu1* alleles were backcrossed to W22 ≥ 4 times. *dcl2-mu1* was isolated from Uniform-Mu line UFMu-12288. All genetic experiments used segregating wild-type progeny as experimental controls. Plants were grown under greenhouse and field conditions.

### Phenotyping and Microscopy

All pollen phenotyping was performed using mature 5mm anthers prior to anthesis.

Individual anthers were suspended in PBS and dissected using forceps and an insulin syringe. Starch viability staining was performed using Lugol solution (Sigma, cat#L6146-1L). Measurements for days to anthesis (DTA) were taken for three replicate crosses (*Tpd1/tpd1; Tpd2/tpd2* x W22) with staggered planting dates in three different field positions. The Leaf Collar Method^107^ was combined with routine manual palpation of the topmost internode to track reproductive stages. Meiotic anthers were dissected, fixed in 4% paraformaldehyde + MBA buffer^108^, and stained with DAPI for visualization. For tetrad viability assays, anthers from the upper floret of an individual spikelet were dissected and stored in MBA. One anther was used for staging and the others were dissected to release the tetrads. Fluorescein diacetate viability staining was performed as previously described^109^. To control for artifacts associated with sample handling, only intact tetrads (4 physically attached spores) were considered.

### Genotyping and marker design

For routine genotyping, tissue discs were collected with a leaf punch and stored in 96 well plates. To extract genomic DNA, 20μl of extraction solution (0.1M NaOH) was added to each well and samples were heated to 95°C for ten minutes then placed immediately on ice. To neutralize this solution, 90μl of dilution solution (10mM Tris + 1mM EDTA, pH to 1.5 with HCl) was added. PCR reactions, using 1-2μl of this solution as template, were performed using GoTaq G2 Green Master Mix (Promega, cat#M7822). Secondary validation of genotyping reactions was performed as needed using the Quick-DNA Plant/Seed Miniprep kit (Zymo Research, cat#D6020). Bulk Illumina and Nanopore data from *Tpd1; Tpd2* seedlings was used for codominant molecular marker design (Supplementary Table 6). When possible, markers based upon simple sequence length polymorphisms (SSLPs) were prioritized, but a number of restriction fragment length polymorphisms (RFLPs) were also designed. W22, *Tpd1/tpd1; Tpd2/tpd2*, and *Tpd1;Tpd2* genomic DNA was used to validate marker segregation prior to use. The *dcl2-mu1* insertion was amplified by combining gene-specific forward and reverse primers with a degenerate terminal inverted repeat (TIR) primer cocktail. The insertion was subsequently validated by Sanger sequencing.

### High Molecular Weight Genomic DNA Extraction

High molecular weight (HMW) genomic DNA was used as input for all Nanopore and bulk Illumina sequencing experiments. For extraction, bulked seedlings were dark treated for 1 week prior to tissue collection. Four grams of frozen tissue was ground under liquid N_2_ and pre-washed twice with 1.0M sorbital. The tissue was then transferred to 20ml pre-warmed lysis buffer (100mM Tris-HCl pH 9.0, 2% w/v CTAB, 1.4M NaCl, 20mM EDTA, 2% PVP-10, 1% 2-mercaptoethanol, 0.1% sarkosyl, 100μg/mL proteinase K), mixed gently, and incubated for 1 hour at 65°C. Organic extraction in phase-lock tubes was performed using 1 vol phenol:chloroform:isoamyl alcohol (25:24:1) followed by 1 vol chloroform:isoamyl alcohol. DNA was precipitated by adding 0.1 vol 3M NaOAc pH 5.2 followed by 0.7 vol isopropanol. HMW DNA was hooked out with a pasteur pipette and washed with 70% EtOH, air dried for 2 minutes, and resuspended in 200μl Tris-Cl pH 8.5 (EB). The solution was treated with 2μl 20mg/ml RNase A at 37°C for 20 minutes followed by 2ul 50mg/ml proteinase K at 50°C for 30 minutes. 194μl EB, 100ul NaCl, and 2μl 0.5M EDTA were added and organic extractions were performed as before. DNA was precipitated with 1.7 vol EtOH, hooked out of solution with a pasteur pipette, washed with 70% EtOH, and resuspended in 50μl EB.

### Nanopore and Hi-C sequencing, TPD genome assembly and annotation

HMW DNA from *Tpd1;Tpd2* BC8S3 was gently sheared by passage through a P1000 pipette 20 times before library preparation with the Oxford Nanopore Technologies Ligation Sequencing gDNA (SQK-LSK109) protocol with the following modifications: 1) DNA repair, end-prep, and ligation incubation times extended to 20 min. each; 2) 0.8x vol. of a custom SPRI bead solution was used for reaction cleanups^110,111^ 3) bead elutions carried out at 50°C for 5 min. Libraries were sequenced on the MinKNOW device with R9.4.1 flow cells. Offline base calling of ONT reads was performed with Guppy 5.0.7 and the R9.4.1 450bps super accuracy model. Reads longer than 1 Kbp were assembled into contigs using Flye 2.9-b1768^45^ with options “--extra-params max_bubble_length=2000000 -m 20000 -t 48 --nano-raw”. The same long reads were aligned to the Flye contigs (filtered to keep only the longest alternatives) using minimap2 2.22-r1101^112^, and these alignments were passed to the PEPPER-Margin-DeepVariant 0.4 pipeline^46^ to polish the initial consensus. To correct remaining SNVs and small indels, two Illumina PCR-free gDNA PE150 libraries were mapped to the long read polished consensus with bwa-mem2 2.2.1^113^ for further polishing with NextPolish 1.3.1^114^ followed by Hapo-G 1.2^115^, both with default options. Two biological replicate samples of BC8S3 leaf tissue were used to prepare Dovetail Omni-C Kit libraries following the manufacturer’s protocol, and sequenced as a PE150 run on a NextSeq500. These Hi-C reads were mapped to the polished contigs with the Juicer pipeline release 1.6 UGER scripts with options “enzyme=none”^116^. The resulting “merged_nodups.txt” alignments were passed to the 3D- DNA pipeline to iteratively order and orient the input contigs and correct misjoins^117^. This initial automatic scaffolding resulted in 11 superscaffolds longer than 10 Mbp. Correcting a single centromeric break during manual review with JBAT^118^ resulted in the expected 10 pseudomolecules. One 6 Mbp contig was identified as bacterial with no contacts, and was discarded. The remaining unscaffolded contigs were of organelle origin (n=9, 625 Kbp), or aligned to the pseudomolecules (n=116, 12 Mbp). Coding gene predictions from the NRGene 2.0 W22 reference^119^ were projected onto the TPD genome assembly using Liftoff 1.6.2^120^ with options “-polish -copies -chroms <CHROM_MAP>”. An average Phred QV score for the assembly was estimated from a 20-mer database of the Illumina reads using merqury 1.4.1^121^ with default options. Assembly completeness was also assessed with BUSCO 5.5.0^122^ with options “-m genome --miniprot”. See Supplementary Table 3 for assembly metrics.

### RNA extraction

Tissue was collected, snap frozen in liquid nitrogen, and stored at -80°C. Samples were ground into a fine powder using a mortar and pestle on liquid nitrogen. 800μl of pre-extraction buffer (100mM Tris-HCl pH 8.0, 150mM LiCl, 50mM EDTA pH 8.0, 1.5% v/v SDS, 1.5% 2- mercaptoethanol) was added and mixed by vortexing. 500μl of acid phenol:chloroform (pH 4.7-5.0) was added and samples were mixed then spun down at 13,000 x g for 15 minutes at 4°C. The aqueous layer was extracted and 1ml Trizol per 200mg input tissue of was added. Samples were mixed by vortex and incubated at RT for 10 minutes. 200μl chloroform per 1ml Trizol was added and samples were mixed by vortexing then incubated at RT for 2 minutes. Samples were then spun down at 13,000 x g for 15 minutes at 4°C. The aqueous phase was extracted and cleaned up using the Zymo RNA Clean and Concentrator-5 kit (Zymo Research, cat#R1013). Only samples with RNA integrity (RIN) scores ≥9 were used for qPCR and sequencing.

### Reverse Transcription and RT-qPCR

For reverse transcription, 1ug of total RNA was treated with ezDNase (ThermoFisher, cat#11766051) according to the manufacturer’s instructions. Reverse transcription (RT) was performed with SuperScript IV VILO Master Mix (ThermoFisher, cat#11756050). Following RT, complementary DNA (cDNA) was diluted 1:20 in dH20 to be used as template in RT-qPCR.

All RT-qPCR experiments were performed on an Applied Biosystems QuantStudio 6 system in 96- well plate format using PowerUp SYBR Green Master Mix (ThermoFisher, cat#A25741). Prior to use in experiments, primer efficiency was tested for each primer set using a standard curve generated from serial dilutions of cDNA template. Only primer sets with efficiencies between 90-110% were used (Supplementary Table 7). For experiments, ≥3 biological replicates (independent cDNA samples from discrete plants) were assayed per genotype, and ≥2 technical replicates were set up for each reaction condition. Raw Ct (cycle threshold) from technical replicates were averaged, and ΔCt (mean Ct^exp^ – mean Ct^ref^) was calculated using *Elfa9* as a housekeeping reference. ΔΔCt values (ΔCt^cond1^ -ΔCt^cond2^) were calculated between genotypes and converted to fold-change (2^(-ΔΔCt)^).

### Whole Genome Sequencing and SNP Calling

HMW DNA from separately maintained *Tpd1;Tpd2* lineages (BC8S3, BC5S2) and bulk segregation analysis (BSA) maternal pools were extracted as detailed above. Libraries were prepared using the Illumina TruSeq DNA PCR-Free kit (Illumina, cat#20015962) with 2μg of DNA input.

Samples were sequenced on a NextSeq500 platform using 2x150bp high output run. Adapter trimming was performed with Cutadapt v3.1^123^. Paired-end reads were aligned to the W22 reference genome^119^ with BWA-MEM v0.7.17^124^. Alignments were filtered by mapping quality (mapQ≥30) and PCR duplicates were removed using Samtools v1.10^125^. SNP calling was performed using Freebayes v1.3.2^126^. Putative SNP calls were filtered by quality, depth, and allele frequency (AF=1) to obtain a high-confidence *mexicana* marker set that was subsequently validated against the *TPD* assembly. For BSA analysis, SNP calls were filtered against the gold-standard *TPD* marker set. Reference and alternate allele frequencies at each marker were calculated and average signal was consolidated into 100kb bins. ΔSNP index was then calculated for each bin in a sliding window.

### Single-pollen grain sequencing

Pollen grains from *Tpd1/tpd1; Tpd2/tpd2* plants were suspended in ice-cold PBS on a microscope slide under a dissecting scope. Individual plump, viable pollen grains were deposited into the 0.2mL wells of a 96-well plate using a p20 pipette. Lysis and whole genome amplification were performed using the REPLI-g single-cell kit (Qiagen, cat#150345) with the following modifications; 1/4^th^ the specified volume of amplification mix was deposited in each well and isothermal amplification was limited to 5 hours. All steps prior to amplification were performed in a UV-decontaminated PCR hood. WGA products were cleaned up using a Genomic DNA Clean & Concentrator kit (Zymo Research, cat#D4067) and yields were quantified using with the QuantiFluor dsDNA system (Promega, cat#E2670) in a 96-well microplate format.

Libraries were prepared using the TruSeq Nano DNA High Throughput kit (Illumina, cat#20015965) with 200ng input. Samples were sequenced on a NextSeq500 platform using 2x101bp high output runs. Quality control, adapter trimming, alignment, and SNP calling were performed as above. BCFtools 1.14^127^ was used to derive genotype calls from single pollen grains at the predefined marker positions and then passed to GLIMPSE 1.1.1^128^ for imputation. All calls at validated marker sites were extracted and encoded in a sparse matrix format (rows = markers, columns = samples) and encoded (1 = alt allele, -1 = ref allele, 0 = missing). To assess *mexicana* introgression in individual pollen grains, mean SNP signal was calculated in 100kb bins across the genome. A sliding window (1Mb window, 200kb step) was applied in order to smooth the data and identify regions with *mexicana* SNP density. To identify genomic intervals overrepresented in surviving *TPD* pollen grains, aggregate allele frequency was calculated across all pollen grains at each marker site.

### RNA sequencing and analysis

Five biological replicates were prepared for each biological condition (*Tpd1/tpd1; Tpd2/tpd2* and *tpd1; tpd2* siblings). 5μg of total RNA was ribosome depleted using the RiboMinus Plant Kit (ThermoFisher, cat#A1083808), and libraries were prepared using the NEXTFLEX Rapid Directional RNA-seq kit (PerkinElmer, cat#NOVA-5138-08). The size distribution of completed libraries was assessed using an Agilent Bioanalyzer, and quantification was performed using a KAPA Library Quantification kit (Roche, cat#KK4824). Libraries were sequenced on a NextSeq500 platform using a 2x150bp high output run. Trimmed reads were aligned to the W22 reference with STAR in two-pass alignment mode^129^. Read counts were assigned to annotated features using featureCounts^130^. For transposable element expression, multi-mapping reads were assigned fractional counts based on the number of identical alignments. Differential expression analysis was performed using edgeR^131^. To avoid false positives, a stringent cutoff (log2FC ≥ 2, FDR ≤ 0.001) was used to call differentially expressed genes. Gene ontology (GO) analysis (Fisher’s exact test, p<0.01) was performed using topGO^132^, and the results were visualized using rrvgo^133^. For data visualization, alignment files were converted to a strand-specific bigwig format using deepTools^134^.

### Small RNA sequencing and analysis

For comparisons between *Tpd1/tpd1; Tpd2/tpd2* and *tpd1; tpd2* pollen, three biological replicates were used. Two biological replicates were used for *dcl2^T^-/-* and *dcl2-mu1-/-* pollen samples. Libraries were constructed with the NEXTFLEX Small RNA-Seq V3 kit (PerkinElmer, cat#NOVA- 5132-06) using 2μg of total RNA input per library and the gel-free size selection protocol. The size distribution of completed libraries was assessed using an Agilent Bioanalyzer, and quantification was performed using a KAPA Library Quantification kit (Roche, cat#KK4824). Libraries were sequenced on a NextSeq500 platform using a 1x76bp run. Adapters were trimmed using cutadapt^123^ and the 4bp UMI sequences on either side of each read were removed.

Reads were filtered using pre-alignment to a maize structural RNA consensus database using bowtie2^135^. Alignment and *de novo* identification of small RNA loci was performed with ShortStack^136^, using a minimum CPM cutoff of 5, and only clusters with clear size bias (21, 22, or 24nt) were retained in downstream analysis. Differential sRNA accumulation was performed with edgeR^131^ (log2FC ≥ 2, FDR ≤ 0.01). The accumulation of size and strand-biased hp-siRNAs was used to identify hairpin loci throughout the genome. For each locus, the underlying primary sequence was tested for reverse complementarity and RNA secondary structure prediction was performed using RNAfold^137^. Non-hairpin siRNA targets were only retained if they showed negligible strand-bias (i.e., evidence of a dsRNA template for processing by a Dicer-like enzyme).

### iPARE sequencing and analysis

For iPARE-seq libraries, 40μg of total RNA was poly(A) selected using a Dynabeads mRNA Purification Kit (ThermoFisher, cat#61006). 1μg of poly(A) RNA was ligated to the 5’ PARE adapter (100pmol) in 10% DMSO, 1mM ATP, 1X T4 RNA ligase 1 buffer (New England Biolabs, cat#B0216L), 25% PEG8000 with 1μl (40U) of RNaseOUT (ThermoFisher, cat#10777019) and 1μl T4 RNA ligase 1 (New England Biolabs, cat#M0204S) in a reaction volume of 100μl. Ligation reactions were performed for 2 hours at 25°C followed by overnight incubation at 16°C. Samples were then purified using RNA Clean XP beads (Beckman Coulter, cat#A63987) and eluted in 18μl dH20. Chemical fragmentation of ligated RNA to ≤200nt was performed using the Magnesium RNA fragmentation kit (New England Biolabs, cat#E6150S). 2ul RNA Fragmentation Buffer was added and samples were incubated at 94°C for 5 minutes followed by a transfer to ice and the addition of 2μl of RNA Stop solution. Samples were purified using the RNA Clean & Concentrator-5 kit (Zymo Research, cat#R1013) and eluted in 11μl H20. Reverse-transcription was performed as follows: 10μl of RNA, 1μl 10mM dNTP, and 2μl random primer mix (New England Biolabs, cat#S1330S) were mixed and incubated for 10 minutes at 23°C then put on ice for 1 minute. The following was then added: 4μl 5X SuperScript IV buffer, 1μl 100mM DTT, 1μl RNaseOUT, and 1μl Superscript IV (200U). The reaction was incubated for 10 minutes at 23°C followed by 10 minutes at 50°C. 80μl of TE was then added to this mixture.

Target indirect capture was performed with 100μl Dynabeads MyOne Streptavidin T1 beads (ThermoFisher, cat#65601) as per manufacturer instructions. 100μl of the RT reaction was used as input, and captured cDNA molecules were eluted in 50μl. Second-strand synthesis was performed using 5U Klenow fragment (New England Biolabs, cat#M0210S) with 100uM dNTPs and 1μM of iPARE adapter primer (5’-NNNNTCTAGAATGCATGGGCCCTCCAAG-3’) for 1 hour at 37°C, incubation at 75°C for 20 minutes. Samples were purified with a 1:1 ratio of AMPure XP SPRI beads (Beckman Coulter – cat#A63880) and resuspended in 51μl EB. 50μl of sample was used for library preparation with the NEB Ultra DNA library kit (New England Biolabs, cat# E7370S). Barcoded samples were sequenced with a NextSeq500 2x150bp high output run. Use of the directional iPARE adapter allows for the retention of directionality even when using a non-directional DNA-seq kit.

Cutadapt^123^ was used to search and recover the adapter sequence in both 5’ and 3’ orientation (forward in read1 or read2 respectively). Read1 adapter reads were trimmed for the 3’ adapter if present, and the 5’ iPARE adapter was subsequently removed. Potential polyA tails were also removed, and only reads ≥20nt were retained. Read2 adapter reads were processed in an identical fashion. Filtered reads were mapped using Bowtie2^135^ and the 5’ position of each read (the cloned 5’ monophosphate) was extracted using BEDtools^138^ with CPM normalization. Small RNA target prediction was performed using psRNATarget^70^.

### Protein Extraction and Western Blotting

Fresh anthers or pollen were collected and snap frozen in liquid nitrogen. Samples were then ground to a fine powder in a mortar and pestle over liquid nitrogen and resuspended in freshly prepared extraction buffer (2mM Tris-HCl pH 7.4, 150mM NaCl, 1mM EDTA, 1% v/v NP40, 5% v/v glycerol, 1mM PMSF, 1ml Roche Protease Inhibitor cocktail per 30g input tissue) and vortexed thoroughly. Samples were then centrifuged at 14,000rpm at 4°C for 5 minutes to pellet cellular debris, and the aqueous fraction was transferred to another tube. This step was then repeated twice more.

Protein extracts were quantified using the Pierce Detergent Compatible Bradford Assay Kit (ThermoFisher, cat#23246) on a Promega Glomax-Multi+ plate reader.

To assess the role of 22nt siRNAs in translational repression, antiserum was raised against a peptide (SRKGAPPSSPPLSPPKLGA) from the Zm00004b012122 protein in collaboration with PhytoAB. Specificity was determined as follows: (1) blots using pollen protein extracts showed a single band at roughly the expected size, and (2) blots using leaf protein extracts showed no band in concordance with expected pollen/anther specificity. A rabbit polyclonal heat shock protein 90-2 (HSP90-2) antibody (Agrisera, cat#AS11 1629), a constitutive isoform with high expression, was used as loading control in all western blot experiments. For comparisons of protein abundance between wild-type and *TPD* pollen/anthers, 2ug of protein was denatured at 95°C for 5 minutes in an appropriate volume of 2X Laemmli buffer (120nM Tris-Cl pH 6.8, 4% v/v SDS, 0.004% bromophenol blue, 20% v/v glycerol, 0.02% w/v bromophenol blue, 350mM DTT). Samples were run on a 4-20% Mini-PROTEAN TGX Precast Gel (BioRad, cat#4561094) with a Precision Plus Protein Dual Xtra Prestained standard (BioRad, cat#1610377).

Transfer to a PVDF membrane was performed using a BioRad Trans-Blot Turbo Transfer system. Membranes were blocked using 5% w/v powdered milk in 1X TBS-T (20 mM Tris, 150mM NaCl, 0.1% Tween-20) for 1 hour at room temperature. Subsequently, the membrane was cut and incubated with primary antibody (1:3,000 dilution in blocking solution) at 4°C overnight with gentle agitation. Three 15-minute membrane washes were performed with 1X TBS-T at room temperature. Membranes were then incubated with a 1:3,000 goat anti-rabbit IgG H&L (PhytoAB, cat#PHY6000) secondary for 1 hour at room temperature. Following three more washes with 1X TBS-T, membranes were incubated for 5 minutes with ECL Prime detection reagent (Amersham, cat#RPN2236) and visualized using a BioRad ChemiDoc Touch Imaging System.

### Esterase Enzymatic Activity Assay

Esterase activity assays were performed using the colorimetric substrate p-Nitrophenyl butyrate (Sigma, cat#N9876) at a final concentration of 1mM in 0.5M HEPES (pH 6.5). For assays using whole 5mm anthers, 100μg of total protein was used as input for each sample, whereas 50μg was used for pollen. Individual samples were prepared in cuvettes at a volume of 1.5mL. Upon addition of the total protein extract, samples were gently mixed and an initial 410nm absorbance reading was taken to serve as a per sample baseline. Samples were then incubated at 30°C and absorbance readings were taken every 5 minutes for a total of 12 timepoints. This experiment was replicated three times for each genotype. All absorbance readings were taken in using a Thermo Scientific Genesys 20 spectrophotometer.

### Detection of selective sweeps in candidate regions associated with *TPD*

We investigated signals of selection in genomic regions associated with *TPD* using selscan v.1.2.0a^139^ to calculate the genome-wide normalized absolute iHS statistics for individual SNPs and in 10kb windows. iHS is suitable for identifying selection in a single population and relies on the presence of ongoing sweeps and a signal of selection from unusually long-range linkage disequilibrium. We also used VCFtools v0.1.16^140^ to calculate Weir & Cockerham’s FST in 10kb windows to assess signals of selection based on changes in allele frequency between populations.

Phased SNPs for modern temperate maize lines, teosinte and *Tripsacum dactyloides* were obtained from Grzybowski et al.^85^ and SNPs for 265 CIMMYT landraces were obtained from Yang et al.^86^ and phased with Beagle v5.4^141^. A phased and imputed set of 42,387,706 genome-wide concatenated SNPs was used for the analysis of selection. The *T*. *dactyloide*s allele was set to be the ancestral allele. A consensus genetic map curated by Ed Coe was obtained from MaizeGDB^142^ and SNP positions were interpolated to genetic positions. Weighted FST was calculated for each unique population pair. For iHS, 10kb windows were binned into 10 quantiles based on the number of SNPs they contained, and empirical *p*-values for each window were calculated within each quantile. The statistic calculated was the number of extreme (top 5%) |iHS| scores per window. Empirical *p*-values for iHS and FST were then calculated from the rank of each window based on the respective statistics. We adjusted these *p*-values for multiple testing of different populations using the Bonferroni method. *TPD*-linked regions (*dcl2*, *rdm1*, *tdr1*, hairpin region) and their 1kb upstream and downstream regions were intersected with the 10kb windows using bedtools v2.30^138^ and assigned the lowest *p*- value of all intersecting windows. To validate our selection scan, we also investigated windows intersecting with a set of four known domestication genes^143^.

## ACKNOWLEDGEMENTS

We thank Kelly Dawe and Jim Birchler for helpful discussions and for sharing cytogenetic data. Research in the Martienssen laboratory is supported by the U.S. National Institutes of Health (NIH) grant R01 GM067014, the National Science Foundation Plant Genome Research Program, and the Howard Hughes Medical Institute. The authors acknowledge assistance from the Cold Spring Harbor Laboratory Shared Resources, which are funded in part by a Cancer Center Support grant (5PP30CA045508). BB was supported by a predoctoral fellowship from the National Science Foundation.

## AUTHOR CONTRIBUTIONS

BB, JK and RAM designed the study; BB, EE, BR, CA and JL performed the experiments; BB, EE, JC, AA, JRI and RAM analyzed the data and/or its significance; BB and RAM wrote the manuscript with contributions from JC, JRI and AA. AS and RAM acquired funding.

## DATA AND MATERIAL AVAILABILITY

Sequencing datasets generated during the current study are available at NCBI (GEO SuperSeries: GSE234925) and datasets used for genome assembly are available at SRA (BioProject: PRJNA937229).

This Whole Genome Shotgun project has been deposited at DDBJ/ENA/GenBank under the accession JARBIH000000000. The version described in this paper is version JARBIH010000000. All materials are available upon request.

## CODE AVAILABILITY

All code is available on Github (https://github.com/martienssenlab/TPD-manuscript).

