## Extended data Figures for "*Teosinte Pollen Drive* guides maize diversification and domestication by RNAi"

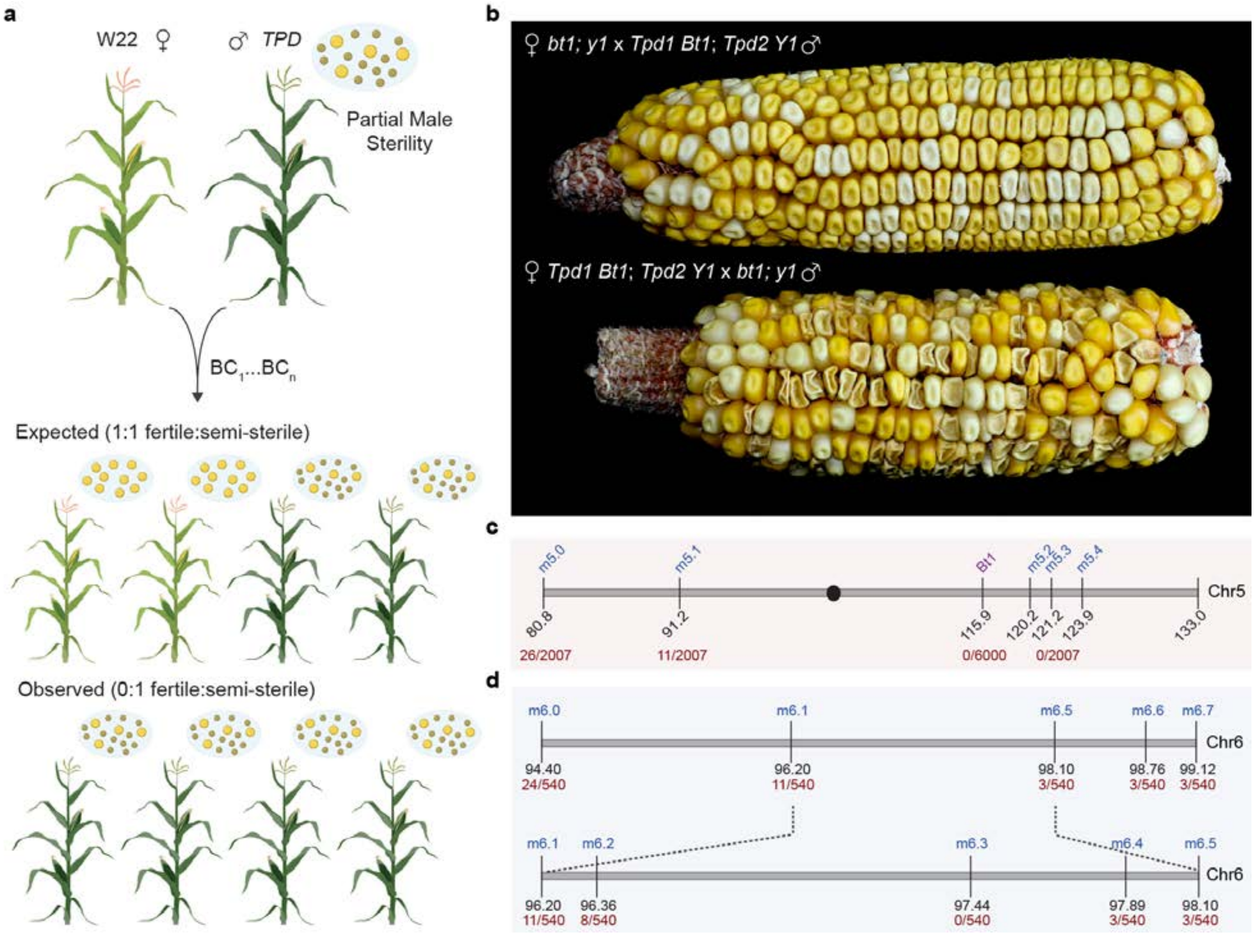


**Extended Data Fig.1: *Teosinte Pollen Drive* and genetic mapping of *Tpd1* and *Tpd2*. a,** Crossing scheme of the *TPD* phenotype. When back-crossed as male, all the progeny of semi-sterile *TPD* plants display the semi-sterile pollen phenotype instead of the expected 1:1 fertile:semi-sterile ratio. **b,** Representative ears from *Tpd1 Bt1/bt1*; *Tpd2 Y1/y1* reciprocal crosses with *bt1; y1* testers, demonstrating severe segregation distortion ("drive" of *Bt1*), but only through the male. *bt1 (brittle1*, collapsed kernels); *y1* (*yellow1*, white kernels). **c,** Summary of molecular and morphological mapping of the *Tpd1* interval. Molecular mapping was performed using *Tpd/++* x W22 segregating progeny, whereas morphological mapping was performed by crossing *Tpd1 Bt1/tpd1 bt1* plants to *bt1* testers. **d,** Molecular mapping of the *Tpd2* interval. SNP markers are shown in blue with recombination frequencies in red.

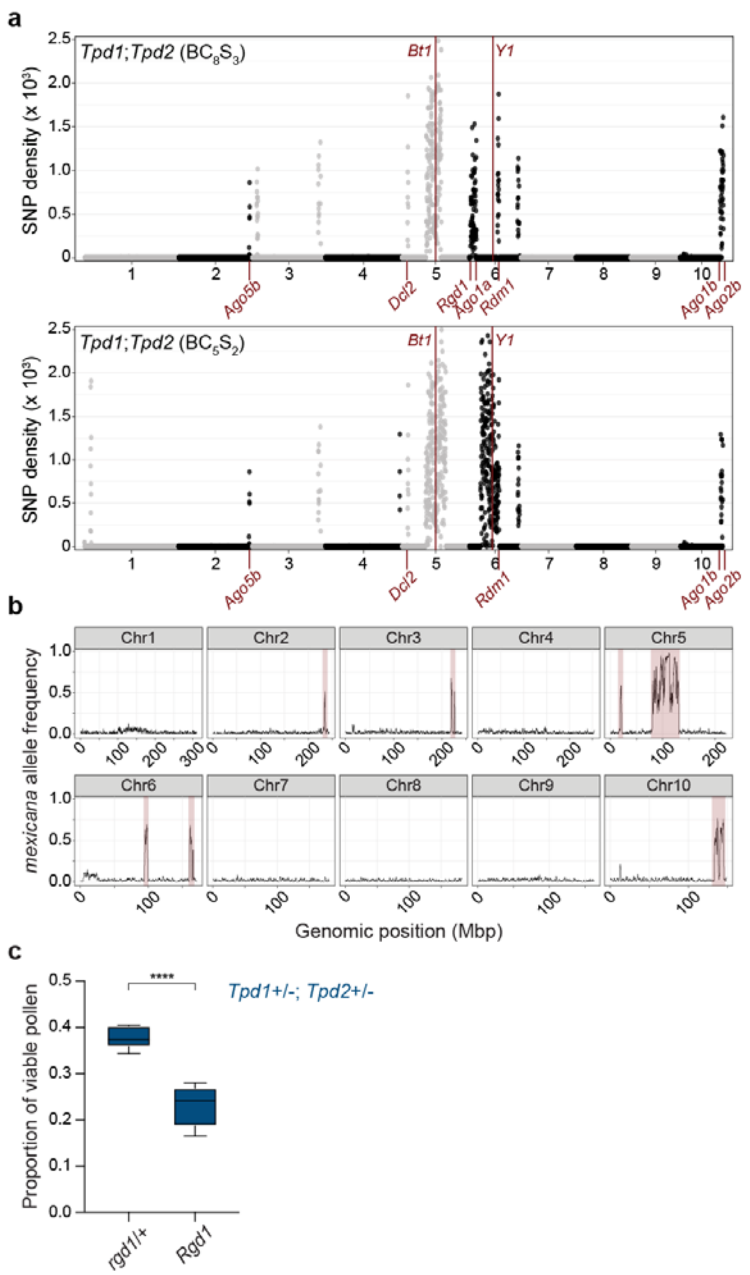


**Extended Data Fig. 2: *mexicana* intervals introgressed into maize carry RNAi genes. a,** Whole genome plots of homozygous *mexicana* SNP density present within *Tpd1; Tpd2* lines. The upper plot corresponds to data from bulked seedlings after 8 backcrosses and 3 self-pollinations ($BC_8S_3$) whereas the lower plot is from $BC_5S_2$ plants. SNP density is consolidated in 250Kb genomic bins. Physical locations for morphological markers *Bt1* and *Y1*, as well as *mexicana* derived RNAi genes, are labeled in red. 7/13 introgression intervals overlap in both independently maintained homozygous lines. **b,** Allele frequency at *mexicana* markers in 96 pollen grains from four different *TPD* plants subjected to single pollen grain sequencing. Regions highlighted in red were over-represented in viable pollen grains. **c,** Quantification of pollen viability in *Tpd1+/-; Tpd2+/-; rgd1/+* and *Tpd1+/-; Tpd2+/-; Rgd1* pollen demonstrating gametophytic suppression via germline segregation of the *rgd1* null allele. n ≥ 9 plants per genotype, ≥ 200 pollen grains per plant. **** $p < 0.0001$ (Welch's t-test).

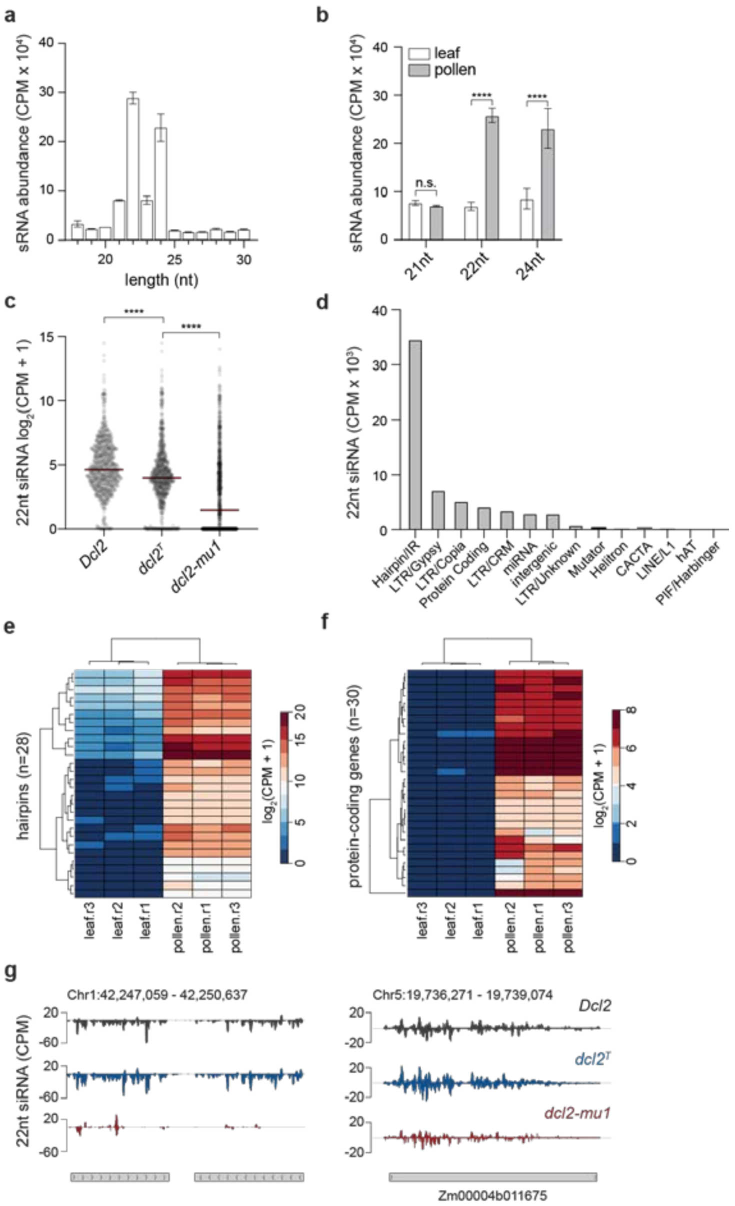


**Extended Data Fig. 3: *DCL2*-dependent 22nt siRNAs from hairpins are prevalent in maize pollen. a,** Distribution and relative abundances of small RNA size classes in WT pollen libraries. Bars indicate mean ± SD. n = 3 biological replicates. **b,** Comparison of relative abundances for 21nt, 22nt, and 24nt sRNA size classes in WT leaf and pollen samples. Both 22nt and 24nt sRNAs show significant increases in pollen. Bars indicate mean ± SD. n = 3 biological replicates. **** p < 0.0001 (Welch's t-test). **c,** Comparison of 22nt sRNA levels in *Dcl2*, *dcl2*$^T$, and *dcl2-mu1* pollen at 804 pollen-specific loci. Values shown are $\log_2$ transformed counts per million (CPM) averaged across replicates. n = 3 replicates per genotype. **** p < 0.0001 (ANOVA test). **d,** Summary of relative contributions for 22nt sRNA producing loci in WT pollen. Hairpin/inverted repeat (IR) hp-siRNAs represent the largest fraction of 22nt species. **e,** Heatmap showing 22nt hp-siRNA levels at hpRNA loci in leaf and pollen. **f,** Heatmap showing 22nt siRNA levels at protein-coding genes in leaf and pollen. **g,** Browser shots showing 22nt hp-siRNA accumulation at a hpRNA locus on chromosome 1 (left) and 22nt siRNA silencing at a representative protein-coding gene. Scale is CPM.

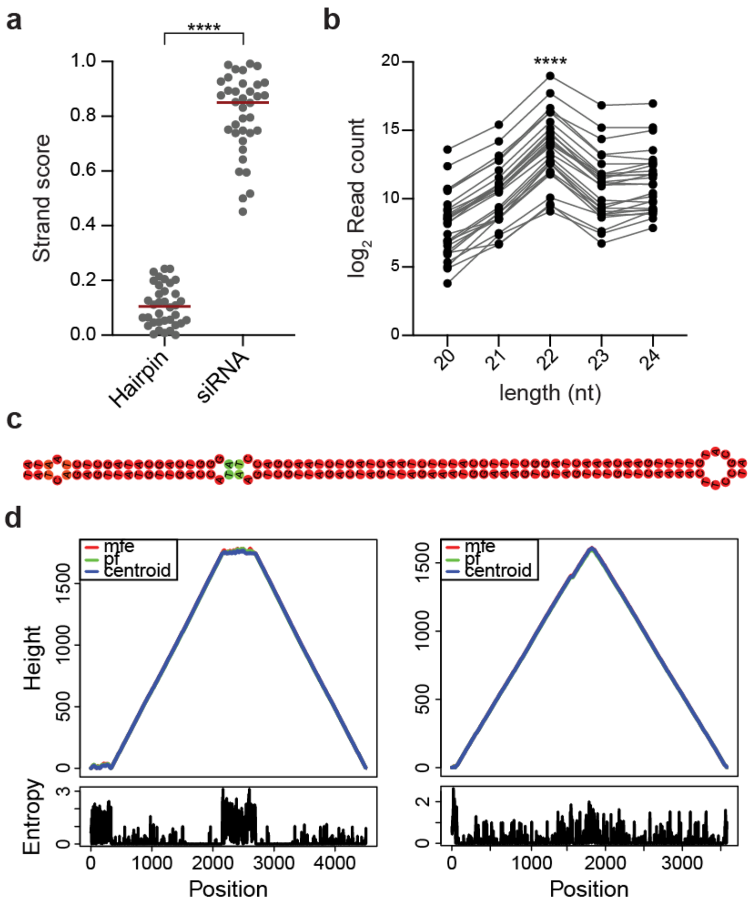


**Extended Data Fig. 4: Validation of highly abundant pollen hairpin precursors. a,** hp-siRNAs are expected to show strand bias. Measurement of strand score (min[plus, minus]/max[plus, minus]) at 28 putative hairpin precursors and randomly selected siRNA clusters from wild-type (W22) maize. A value of zero indicates complete strand bias, whereas a value of 1 indicates unbiased accumulation from both strands. n = 28. **** $p < 0.0001$ (Welch's t-test). **b,** log2 read count at hairpin precursors indicate 22nt size bias. n = 28. **** $p < 0.0001$ (ANOVA). **c,** Example of a 73nt stretch from a 4,480bp hairpin precursor demonstrating near-complete reverse complementarity. **d,** Mountain plots measuring thermodynamic stability of the *Tpd1* hairpin from *mexicana* and another randomly selected hairpin structure.

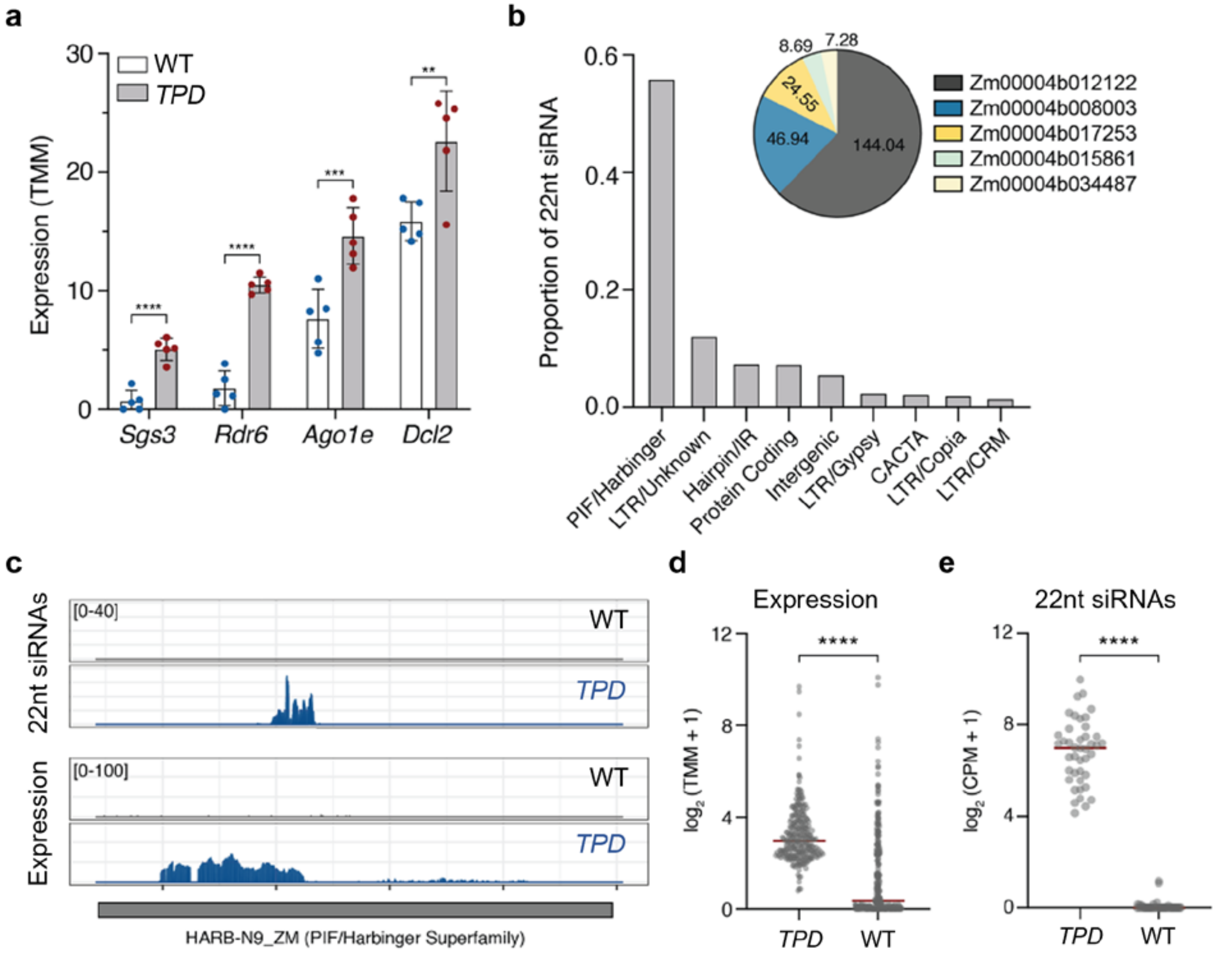


**Extended Data Fig. 5: Origins and targets of 22nt small RNAs in *TPD* Pollen. a,** RNAi genes (*Sgs3*/*Rgd1*, *Rdr6*, *Ago1e* and *Dcl2*) associated with 22nt biogenesis and function are upregulated in *TPD* pollen. Expression is shown in TMM normalized counts. Bars show mean ± SD. n = 5 replicates per condition. **** $p < 0.0001$, *** $p < 0.001$ , ** $p < 0.01$ (FDR). **b,** Relative abundances of *TPD*-dependent 22nt siRNAs mapping to annotated elements. Pie chart inset shows proportions of 22nt siRNAs targeting genes in CPM. **c,** Browser shot showing transcriptional activation at PIF/Harbinger elements in *TPD* pollen as well as 22nt siRNA accumulation. **d,** Quantification of mRNA expression at 258 PIF/Harbinger superfamily elements in WT and *TPD* pollen. **** $p < 0.0001$ (Mann-Whitney test). **e,** 22nt siRNA levels at 42 PIF/Harbinger elements in WT and *TPD* pollen. **** $p < 0.0001$ (Mann-Whitney test).

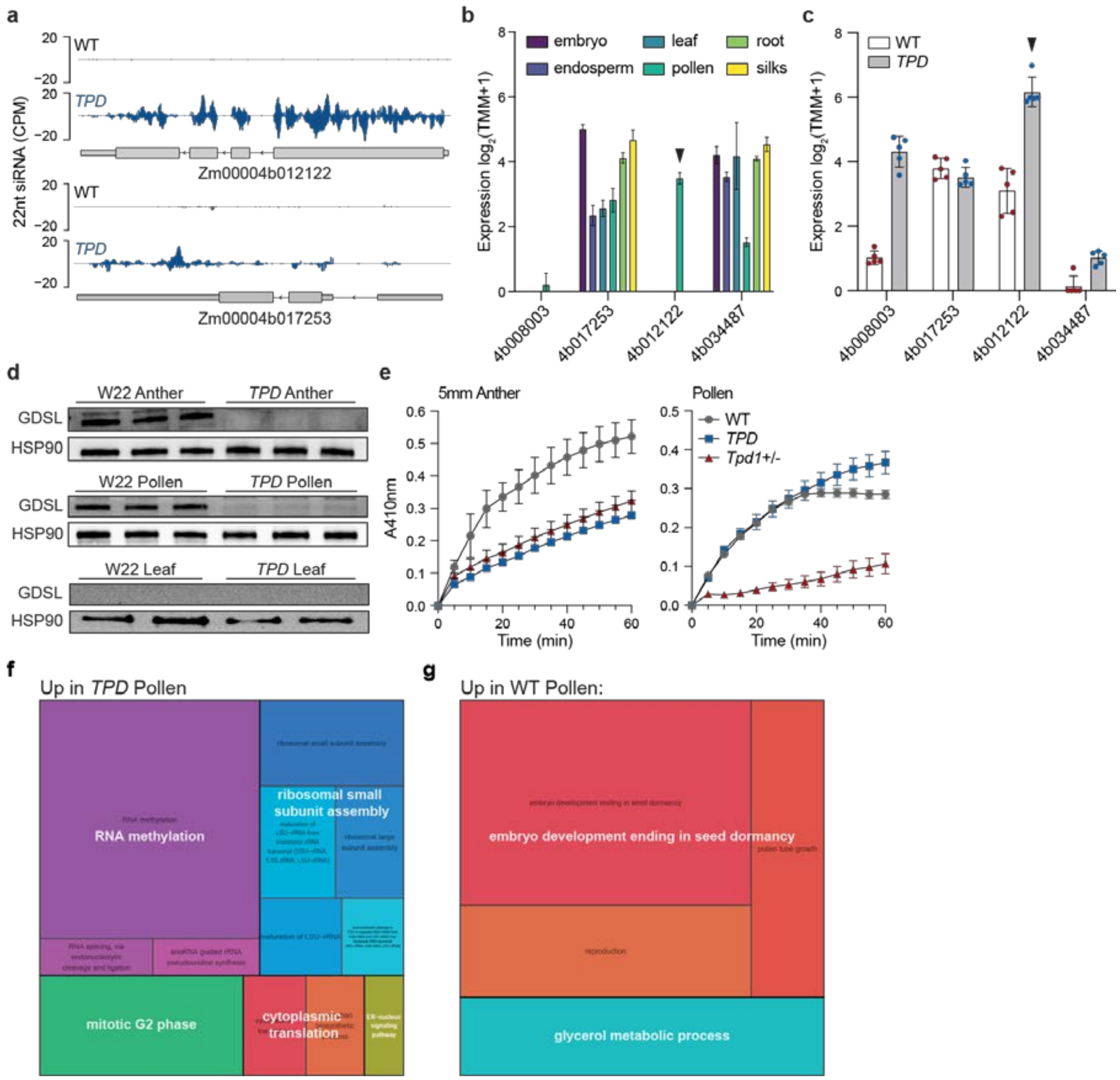


**Extended Data Fig. 6: *TPD*-dependent silencing of a GDSL lipase disrupts lipid metabolism. a,** Browser shots showing ectopic accumulation of 22nt siRNA at protein-coding genes in *TPD* pollen. Scale in counts per million (CPM). **b,** RNA-seq tissue expression of 22nt siRNA targets specific to *TPD* pollen, data from Walley et al. 2016[144]. Bars show mean ± SD. n = 3 replicates per tissue. **c,** RNA-seq expression of 22nt siRNA targets in WT and *TPD* pollen. Bars show mean ± SD. n = 5 replicates per condition. **d,** Western blot comparing TDR1 protein levels in WT and *TPD* pollen, anthers, and leaf. Protein levels were normalized using Heat Shock Protein 90 (HSP90). **e,** p-nitrophenyl butyrate esterase activity assay in 5mm anthers and pollen from WT, *TPD*, and *Tpd1+/-* plants. **f, g,** GO term biological processes up-regulated in **f,** *TPD* and **g,** WT pollen (FDR ≤ 0.001). Upregulated genes in *TPD* pollen were associated with RNA metabolism, ribosome assembly, and cytoplasmic translation as well as G2 mitotic arrest. This could reflect translational repression via 22nt siRNAs. Interestingly, a subset of genes associated with endoplasmic reticulum (ER)-nucleus signaling was also up-regulated[145], while genes associated with glycerol metabolism, the primary backbone for TAG synthesis, were downregulated. In pollen, the accumulation of TAGs in lipid droplets (LDs) is critical for proper membrane expansion and pollen tube growth[146].

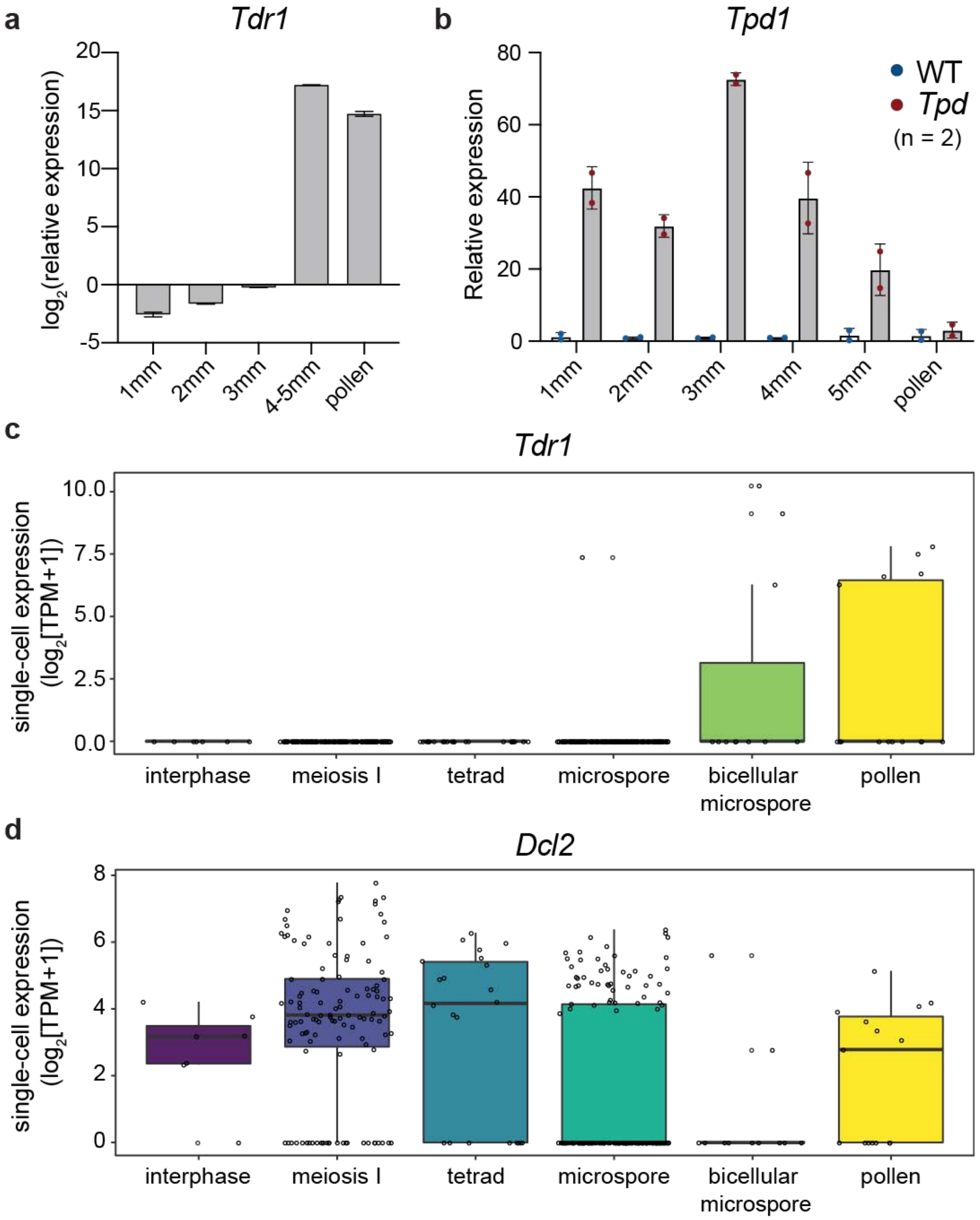


**Extended Data Fig. 7: *Tpd1* and *Dcl2* are expressed pre-meiotically, whereas *Tdr1* is expressed in microspore and pollen. a,** RT-qPCR of the *Tdr1* transcript throughout anther development and in mature pollen. Bars show mean ± SD. n = 2 replicates per condition. **b,** RT-qPCR of the *Tpd1* transcript during anther development and in mature pollen in WT and *Tpd*. Bars show mean ± SD. n = 2 replicates per condition. **c, d,** Single-cell expression at different stages of meiosis of **c,** *Tdr1* and **d,** *Dcl2*, using single cell RNAseq data[73]. Early and late expression of *Dcl2* coincides with *Tpd1* and *Tdr1*, respectively.

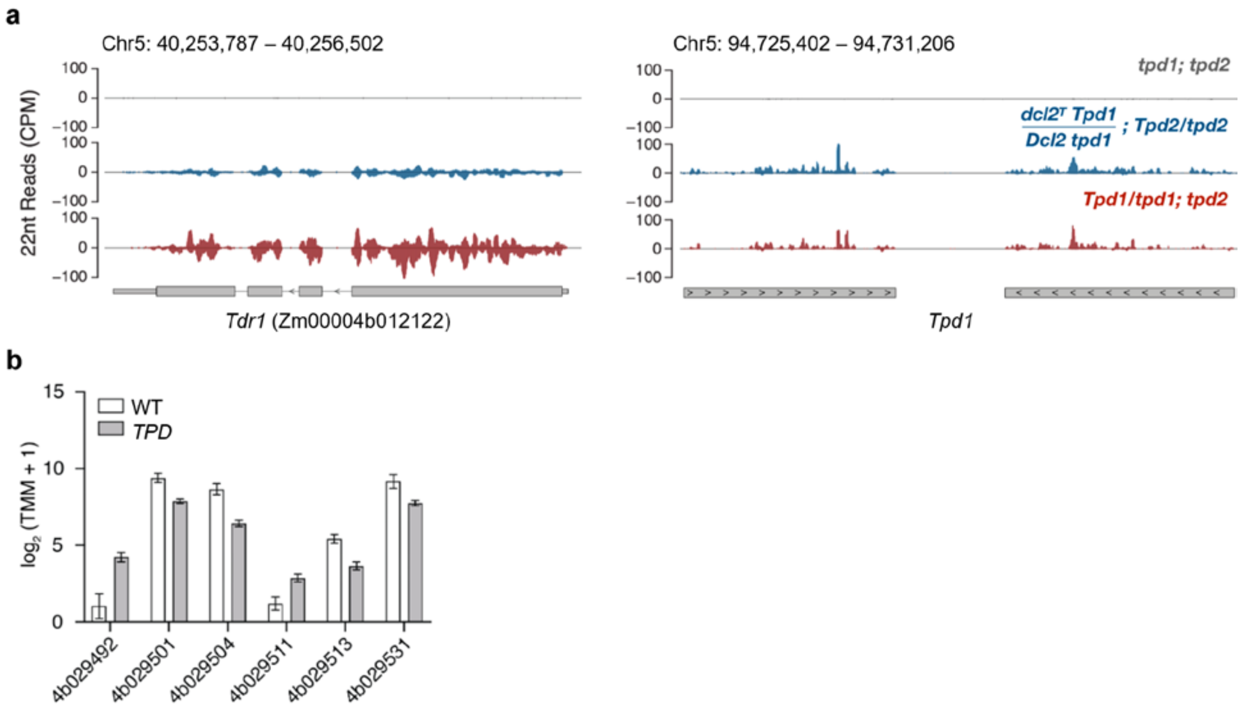


**Extended Data Fig 8: *Tpd2* suppresses 22nt secondary small RNAs. a.** Browser shots showing ectopic accumulation of 22nt siRNAs at *Tdr1* (left) and *Tpd1* hairpin (right) in fertile (*tpd1; tpd2*, grey), drive (*Dcl2$^T$ Tpd1/Dcl2 tpd1; Tpd2/tpd2*, blue) and sterile (*Dcl2 Tpd1/Dcl2 tpd1; tpd2*, red) pollen from maternal segregants. Scale in counts per million (CPM). *Tpd2* and *Dcl2$^T$* reduce small RNAs from *Tdr1* (left) but not from the *Tpd1* hairpin (right), consistent with a cell autonomous role in secondary small RNA biogenesis and silencing. **b.** *Rdm1* (Zm00004b029511) is one of six genes in the *Tpd2* interval expressed in pollen, and is overexpressed in *TPD* pollen.

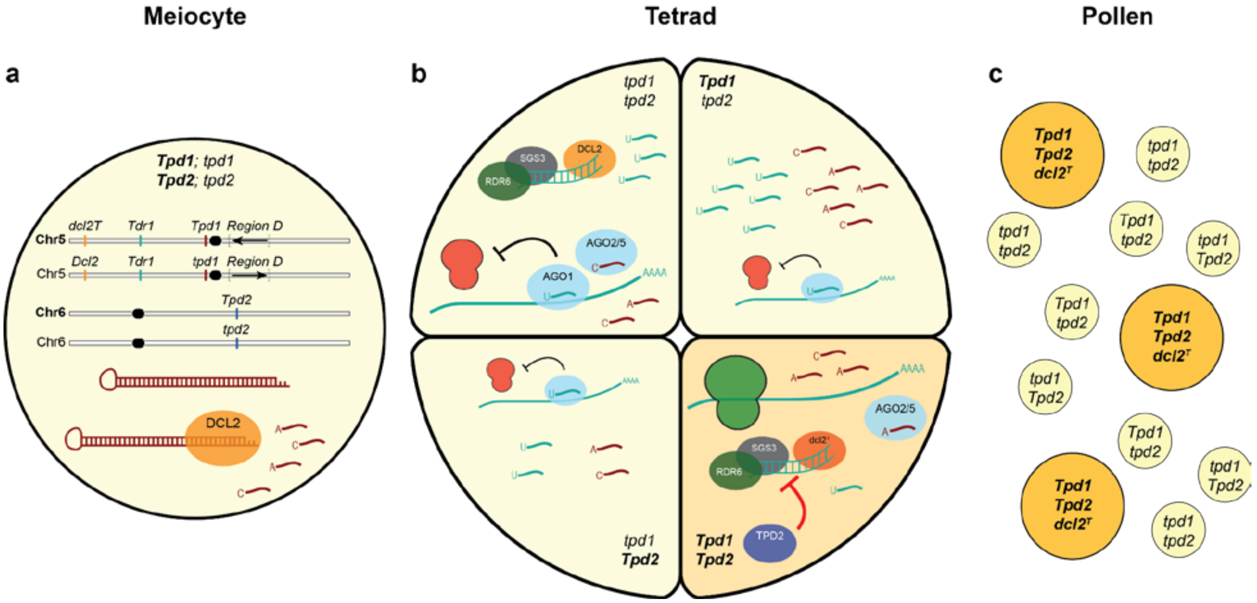


**Extended Data Fig. 9: Mechanistic model of *Teosinte Pollen Drive*. a,** The *TPD* system is defined by *mexicana* introgression intervals on chromosomes 5 and 6. *Tpd1* encodes a pre-meiotically expressed *mexicana*-specific hairpin that produces abundant 22nt hp-siRNAs. **b**, These hp-siRNAs trigger secondary siRNAs amplification by RDR6 and SGS3/RGD1 at the *Tdr1* gene when it starts being transcribed at the late tetrad stage, which in turn target *Tdr1* for translational repression (red ribosomes). In surviving microspores (dark yellow background) *Tpd2* and *dcl2$^T$* repress secondary siRNAs processing, restoring translation and fertility (green ribosome). **c**, Only pollen grains of the genotype *dcl2$^T$ Tpd1*; *Tpd2* are viable, and all other competing gametes are eliminated. Other RNAi genes (*Sgs3*/*Rgd1*, *Ago1*, *Ago2*, *Ago5*) can act as partial suppressors by affecting levels of siRNAs.

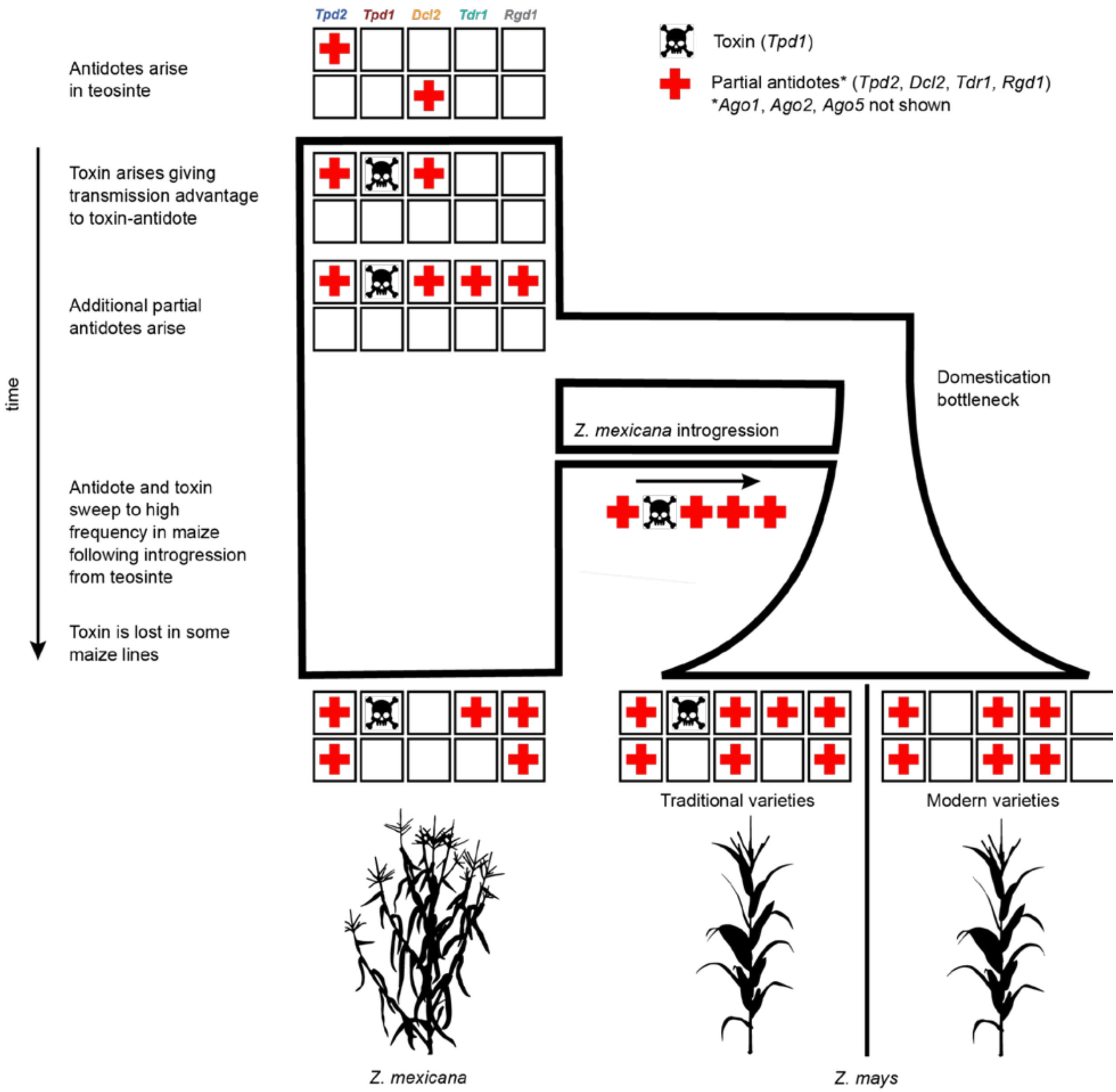


**Extended Data Fig. 10: Evolutionary model of *Teosinte Pollen Drive*.** After the antidotes arise in an ancestral teosinte population, the *Tpd1* toxin arises and gains a transmission advantage when linked to the antidote genes. In extant populations of *Z. mexicana* and *Z. mays*, some antidotes are fixed, while others are polymorphic or lost. The demographic model was based on Beissinger et al. 2016[147] and the conceptual framework of selfish evolution was adapted from Sweigart et al. 2019[148]. The silhouette of *Z. mexicana* was adapted from Iltis 1983[149] and that of *Z. mays* was obtained from phylopic.org.
