## Supplementary Tables1-7 for "*Teosinte Pollen Drive* guides maize diversification and domestication by RNAi"

**Supplementary Table 1: Phenotypic Segregation Ratios for Reciprocal *TPD* Crosses**

| Cross | Fertile | Semi-Sterile | Sterile | Chi-square |
|---|---|---|---|---|
| *TPD* ♂ | 0 | 142 | 0 | $\chi^2$(2:1:1) = 164.29, p < 0.00001 |
| *TPD* ♂ | 0 | 179 | 0 | $\chi^2$(2:1:1) = 209.29, p < 0.00001 |
| *TPD* ♂ | 0 | 219 | 0 | $\chi^2$(2:1:1) = 257.27, p < 0.00001 |
| *TPD* ♀ | 144 | 89 | 58 | $\chi^2$(2:1:1) = 3.301, p = 0.192 |
| *TPD* ♀ | 159 | 109 | 71 | $\chi^2$(2:1:1) = 5.234, p = 0.073 |
| *TPD* ♀ | 106 | 60 | 54 | $\chi^2$(2:1:1) = 0.193, p = 0.908 |

**Supplementary Table 2: Genotypic Segregation Ratios for Reciprocal *TPD* Crosses**

| Parent | *tpd1; tpd2* | *Tpd2+/-* | *Tpd/++* | *Tpd1+/-* | Chi-square |
|---|---|---|---|---|---|
| *TPD* ♂ | 0 | 0 | 142 | 0 | $\chi^2$(1:1:1:1) = 162.44, p < 0.00001 |
| *TPD* ♂ | 0 | 0 | 179 | 0 | $\chi^2$(1:1:1:1) = 204.41, p < 0.00001 |
| *TPD* ♂ | 0 | 0 | 219 | 0 | $\chi^2$(1:1:1:1) = 254.37, p < 0.00001 |
| *TPD* ♀ | 81 | 63 | 89 | 58 | $\chi^2$(1:1:1:1) = 4.447, p = 0.217 |
| *TPD* ♀ | 69 | 90 | 109 | 71 | $\chi^2$(1:1:1:1) = 6.029, p = 0.110 |
| *TPD* ♀ | 51 | 55 | 60 | 54 | $\chi^2$(1:1:1:1) = 0.378, p = 0.945 |

**Supplementary Table 3: Genome Assembly Metrics.** Values calculated on scaffolded assemblies with contigs split by 10 or more consecutive Ns. Repeat content determined by RepeatMasker with the combined MTEC repeat library “maizeTE02052020” distributed at https://github.com/oushujun/MTEC.

| | **TPD**<br>**TPD 1.0 2021** | **W22**<br>**W22 Reference NRGENE-2.0 2018** |
|---|---|---|
| **Contigs** | | |
| # | 945 | 49,111 |
| Longest (Kbp) | 44,353 | 831 |
| N50 (Kbp) | 7,744 | 88 |
| **Scaffolds** | | |
| # > 10Mbp | 10 | 10 |
| Span (Mbp) | 2,112 | 2,134 |
| Longest (Mbp) | 306 | 311 |
| Gap bases (%) | 0.02 | 1.90 |
| **Annotations** | | |
| Coding Genes (W22 liftover) | 40,564 | 40,961 |
| Interspersed Repeats (%) | 81.37 | 80.04 |
| CentC masked (Kbp) | 2,876 | 137 |
| **QC** | | |
| merqury QV estimate | 34.2 | - |
| Complete BUSCOs (%) | 90.6 | 91.1 |
| Missing BUSCOs (%) | 0.3 | 0.6 |

**Supplementary Table 4: Empirical *p*-values for selection scan with a windowed weighted $F_{ST}$ statistic for *Teosinte Pollen Drive*-linked (*TPD*-linked) regions and a validation set of domestication genes.** See Supplementary Table 5 for gene coordinates. Empirical *p*-values for each gene are shown for the overlapping window with the lowest *p*-value. Significant *p*-values <0.05 are indicated with an asterisk for both unadjusted and Bonferroni-adjusted values accounting for each population pair tested.

| Group | Gene | ***mexicana*** **- Landrace** | ***parviglumis*** **- Landrace** | **Landrace - Modern** | ***mexicana*** **- Modern** | ***parviglumis*** **- Modern** |
|---|---|---|---|---|---|---|
| | | **Windowed $F_{ST}$ empirical *p*-values** | | | | |
| *TPD*-linked | *dcl2* | 0.321 | 0.584 | 0.479 | 0.132 | 0.345 |
| *TPD*-linked | hairpin | 0.305 | 0.503 | 0.742 | 0.106 | 0.255 |
| *TPD*-linked | *rdm1* | 0.510 | 0.660 | 0.326 | 0.585 | 0.802 |
| *TPD*-linked | *tdr1* | 0.263 | 0.324 | 0.166 | 0.096 | 0.124 |
| Domestication | *gt1* | 0.128 | 0.058 | 0.020* | 0.213 | 0.164 |
| Domestication | *tb1* | 0.016* | 0.110 | 0.329 | 0.004* | 0.047* |
| Domestication | *tga1* | 0.032* | 0.003* | 0.356 | 0.021* | 0.004* |
| Domestication | *zagl1* | 0.025* | 0.001* | 0.201 | 0.003* | 0.001* |
| | | **Windowed $F_{ST}$ empirical adjusted *p*-values** | | | | |
| *TPD*-linked | *dcl2* | 1.000 | 1.000 | 1.000 | 0.662 | 1.000 |
| *TPD*-linked | hairpin | 1.000 | 1.000 | 1.000 | 0.529 | 1.000 |
| *TPD*-linked | *rdm1* | 1.000 | 1.000 | 1.000 | 1.000 | 1.000 |
| *TPD*-linked | *tdr1* | 1.000 | 1.000 | 0.828 | 0.482 | 0.622 |
| Domestication | *gt1* | 0.638 | 0.290 | 0.100 | 1.000 | 0.822 |
| Domestication | *tb1* | 0.082 | 0.551 | 1.000 | 0.021* | 0.236 |
| Domestication | *tga1* | 0.158 | 0.016* | 1.000 | 0.105 | 0.020* |
| Domestication | *zagl1* | 0.123 | 0.004* | 1.000 | 0.013* | 0.003* |

**Supplementary Table 5: Selection scan with a windowed |iHS| statistic in *Teosinte Pollen Drive*-linked (*TPD*-linked) regions and a validation set of domestication genes in teosinte and maize populations.** Counts of individual significant ($p$<0.05) SNPs as well as empirical $p$-values for 10kb windows are shown. Genomic coordinates for *dcl2* (Zm00001eb219690), *tdr1* (Zm00001eb224090) and *rdm1* (Zm00001eb275620) are for the public maize B73 NAMv5 reference genome and annotation from MaizeGDB. Coordinates for the domestion genes are based on the same assembly and annotation. Significant $p$-values <0.05 are indicated with an asterisk. SNP counts and per window $p$-values are shown as unadjusted and Bonferroni-adjusted values accounting for each population tested. Cases in which there was insufficient data to calculate a $p$-value are shown as "NA".

| Group | Gene | Genomic coordinates | ***mexicana*** | ***parviglumis*** | **Landrace** | **Modern** |
|---|---|---|---|---|---|---|
| | | | **Count of \|iHS\| significant SNPs** | | | |
| *TPD*-linked | *dcl2* | Chr5:20,831,070-20,848,118 (-) | 46 | 6 | 1 | 18 |
| *TPD*-linked | *tdr1* | Chr5:40,759,118-40,761,756 (-) | 1 | 0 | 0 | 0 |
| *TPD*-linked | hairpin | Chr5:95,990,889-96,002,706 (+) | 4 | 30 | 1 | 0 |
| *TPD*-linked | *rdm1* | Chr6:107,817,218-107,820,394 (+) | 0 | 2 | 0 | 14 |
| Domestication | *zagl1* | Chr5:20,831,070-20,848,118 (-) | 48 | 40 | 0 | 0 |
| Domestication | *gt1* | Chr5:40,759,118-40,761,756 (-) | 0 | 0 | 0 | 3 |
| Domestication | *tb1* | Chr5:95,990,889-96,002,706 (+) | 5 | 2 | 2 | 0 |
| Domestication | *tga1* | chr1:272330564-272332648 (+) | 2 | 0 | 0 | 0 |
| | | | **Count of \|iHS\| adjusted significant SNPs** | | | |
| *TPD*-linked | *dcl2* | Chr5:20,831,070-20,848,118 (-) | 10 | 0 | 0 | 0 |
| *TPD*-linked | *tdr1* | Chr5:40,759,118-40,761,756 (-) | 0 | 0 | 0 | 0 |
| *TPD*-linked | hairpin | Chr5:95,990,889-96,002,706 (+) | 0 | 10 | 0 | 0 |
| *TPD*-linked | *rdm1* | Chr6:107,817,218-107,820,394 (+) | 0 | 0 | 0 | 6 |
| Domestication | *zagl1* | Chr1:4932248-4948340 (-) | 0 | 12 | 0 | 0 |
| Domestication | *gt1* | chr1:23433554-23435122 (+) | 0 | 0 | 0 | 3 |
| Domestication | *tb1* | chr1:272330564-272332648 (+) | 4 | 1 | 2 | 0 |
| Domestication | *tga1* | chr4:46647932-46652896 (+) | 0 | 0 | 0 | 0 |
| | | | **Windowed \|iHS\| empirical *p*-values** | | | |
| *TPD*-linked | *dcl2* | Chr5:20,831,070-20,848,118 (-) | 0.008* | 0.341 | 0.316 | 0.079 |

| | | | | | | |
|---|---|---|---|---|---|---|
| *TPD*-linked | *tdr1* | Chr5:40,759,118-40,761,756 (-) | 0.377 | 0.649 | 0.759 | 0.468 |
| *TPD*-linked | hairpin | Chr5:95,990,889-96,002,706 (+) | 0.216 | 0.171 | 0.493 | 0.215 |
| *TPD*-linked | *rdm1* | Chr6:107,817,218-107,820,394 (+) | 0.421 | 0.414 | 0.737 | 0.021* |
| Domestication | *zagl1* | Chr5:20,831,070-20,848,118 (-) | 0.020* | 0.039* | NA | NA |
| Domestication | *gt1* | Chr5:40,759,118-40,761,756 (-) | 0.913 | 0.373 | 0.780 | 0.410 |
| Domestication | *tb1* | Chr5:95,990,889-96,002,706 (+) | 0.021* | 0.116 | 0.054 | 0.188 |
| Domestication | *tga1* | chr1:272330564-272332648 (+) | 0.315 | 0.477 | 0.641 | 0.673 |
| | | | **Windowed \|iHS\| empirical adjusted *p*-values** | | | |
| *TPD*-linked | *dcl2* | Chr5:20,831,070-20,848,118 (-) | 0.031* | 1.000 | 1.000 | 0.314 |
| *TPD*-linked | *tdr1* | Chr5:40,759,118-40,761,756 (-) | 1.000 | 1.000 | 1.000 | 1.000 |
| *TPD*-linked | hairpin | Chr5:95,990,889-96,002,706 (+) | 0.863 | 0.686 | 1.000 | 0.861 |
| *TPD*-linked | *rdm1* | Chr6:107,817,218-107,820,394 (+) | 1.000 | 1.000 | 1.000 | 0.085 |
| Domestication | *zagl1* | Chr1:4932248-4948340 (-) | 0.080 | 0.156 | NA | NA |
| Domestication | *gt1* | chr1:23433554-23435122 (+) | 1.000 | 1.000 | 1.000 | 1.000 |
| Domestication | *tb1* | chr1:272330564-272332648 (+) | 0.082 | 0.465 | 0.217 | 0.753 |
| Domestication | *tga1* | chr4:46647932-46652896 (+) | 1.000 | 1.000 | 1.000 | 1.000 |

**Supplementary Table 6: Genotyping markers used in this study.** The “type” column refers to the general marker design associated with the primer sequence. RFLP markers will denote the associated restriction enzyme in the “type” column.

| Name | Type | Sequence (5’-3’) | WT | Mut/*TPD* |
|---|---|---|---|---|
| *dcl2*$^{T}$f | BstNI | AGCGCCATTTACAATTTCAGCA | 183, 211 | 394 |
| *dcl2*$^{T}$-r | BstNI | TGTTGCCAGTGAATCAGCACTA | 183, 211 | 394 |
| m5.0-f | SSLP | TTAGTAGTGTCCTGGCGCTC | 295 | 165 |
| m5.0-r | SSLP | GTGAGGGACTAGGGCATGTG | 295 | 165 |
| m5.1-f | SSLP | CCTGCATAGAGATGCCATCAA | 310 | 180 |
| m5.1-r | SSLP | CGACGACGACTCATCCACGA | 310 | 180 |
| m5.2-f | SSLP | TGTCTTCCTCCAACTGTGCT | 312 | 211 |
| m5.2-r | SSLP | ACTGCCCAAAGAGCATGTGT | 312 | 211 |
| m5.3-f | MfeI | AATGGTGTTTTCCTTGGCATTCA | 253, 249 | 502 |
| m5.3-r | MfeI | GTACCATGCACTCATCCCGAA | 253, 249 | 502 |
| m5.4-f | TseI | GGATCATGGAGTGCGTGCAG | 450 | 161, 289 |
| m5.4-r | TseI | AACCAGCGCTCCTCAAAGTT | 450 | 161, 289 |
| m6.0-f | SSLP | ACTGAGTAACCAATGCCAGA | 445 | 363 |
| m6.0-r | SSLP | GCAGCCTTCCAGTTCCTGTA | 445 | 363 |
| m6.1-f | SSLP | GATCTACTTGCACGAGAGCACC | 495 | 374 |
| m6.1-r | SSLP | CGGAGTAATTTCCTTGGGACA | 495 | 374 |
| m6.2-f | AflIII | CTACCAACTGCTCCTGAGATGG | 116, 96 | 212 |
| m6.2-r | AflIII | CGTTGACGAATATTGATTGTAGCCA | 116, 96 | 212 |
| m6.3-f | NdeI | TTGCTCCAACCTTGTCACCT | 124, 100 | 224 |
| m6.3-r | NdeI | GCTATCCGCAAACAGCGAGA | 124, 100 | 224 |
| m6.4-f | BccI | TCCTTCTCCTCTTCCCTCGG | 250 | 110, 140 |
| m6.4-r | BccI | AGGAACCCTGTTTGACGATCT | 250 | 110, 140 |
| m6.5-f | BglIII | TCCACAGAAGGACAGCAAAAGGA | 174, 65 | 239 |
| m6.5-r | BglIII | TAGGGTTTGTTGTGGCTGCT | 174, 65 | 239 |
| m6.6-f | PvuII | GGCCAAGTTGTTCAAGAAGCAT | 165, 85 | 254 |
| m6.6-r | PvuII | GCGTGGCCCCTTCTCTTATT | 165, 85 | 254 |
| m6.7-f | NdeI | GGGCATCGTGTCATTGAAGG | 231 | 115, 116 |
| m6.7-r | NdeI | TGCAACCTCTCAGGTCTAAG | 231 | 115, 116 |
| *lbl-rgd1*-f | MwoI | GCCCATCTGGATCTGAAGTC | 262, 227 | 489 |
| *lbl-rgd1*-r | MwoI | TTGGTGGCCACACTATCTCA | 262, 227 | 489 |
| *dcl2mu1*-f | Mu insertion | GTGTCCGCGTTCCAGAAGTC | 906 | 800 |
| *dcl2mu1*-r | Mu insertion | TAAAGGTTGTCCATTGGGCGTT | 906 | 800 |
| TIR4 | TIR | GCCTCCATTTCGTCGAATCCC | - | - |
| TIR6 | TIR | GCCTCTATTTCGTCGAATCCG | - | - |
| Gdsl-f | HiII check | GAGCGTCTCCTTCAACCCAA | 983 | - |
| Gdsl-r | HiII check | TCCTACTCCTCAGTTGGGGG | 983 | - |
| Dcl2-f | HiII check | GGCCTAGAATTCAGTTGCGG | 717 | - |
| Dcl2-r | HiII check | GAACACGCTTGTGTTCCTCG | 717 | - |

**Supplementary Table 7: RT-qPCR primers used in this study.** All RT-qPCR primers used in this study were designed to bridge exon junctions (if present) in order to increase specificity. Expected amplicon sizes corresponding to a cDNA or gDNA template are listed.

| Name | Target | Sequence | cDNA | gDNA |
|---|---|---|---|---|
| *Dcl2q-1f* | *Dcl2* – exon 1,2 | CCCAAAAGGACACACAGCTTTC | 79 | 159 |
| *Dcl2q-1r* | *Dcl2* – exon 1,2 | GCATATCTGATCTTGGAGTATGGC | 79 | 159 |
| *Dcl2q-2f* | *Dcl2* – exon 5,6 | TCGACAAAAACAATGCATCTCAAAT | 104 | 658 |
| *Dcl2q-2r* | *Dcl2* – exon 5,6 | GCTGAGGATCTGCAAGATGG | 104 | 658 |
| *Dcl2q-3f* | *Dcl2* – exon 18,19 | CATTCTGAAGGGTGTCGGGT | 174 | 417 |
| *Dcl2q-3r* | *Dcl2* – exon 18,19 | GGTCCATCACATGGTTGGGAA | 174 | 417 |
| *Gdslq-1f* | *Gdsl* – exon 1,2 | GAGCGTCTCCTTCAACCCAA | 81 | 249 |
| *Gdslq-1r* | *Gdsl* – exon 1,2 | ACAAGGCTACTGGCAGGTTC | 81 | 249 |
| *Gdslq-2f* | *Gdsl* – exon 2,3 | CCCACCTTCGCCTTGTACTC | 153 | 250 |
| *Gdslq-2r* | *Gdsl* – exon 2,3 | CCTGAACAAGAGCCTCACCC | 153 | 250 |
| *Elfa9q-1f* | *Efla9* – exon 1,2 | CAAGCTGACTGTGCTGTTCTTA | 181 | 928 |
| *Elfa9q-1r* | *Efla9* – exon 1,2 | GCCTTTGAATACTTGGGTGTAGTAG | 181 | 928 |
| *Elfa9q-2f* | *Efla9* – exon 1,2 | CCAAGCTGACTGTGCTGTTCTT | 198 | 945 |
| *Elfa9q-2r* | *Efla9* – exon 1,2 | AATCTCATCATAACGGGCCTTTGAA | 198 | 945 |
| *Rdr6q-1f* | *Rdr6* – no exons | TGTTTTGCTTCAGCTGGGGA | 128 | 128 |
| *Rdr6q-1r* | *Rdr6* – no exons | CACGCGAATGAAGCATCTGG | 128 | 128 |
| *Rdr6q-2f* | *Rdr6* – no exons | GCCTCTCACCTTTCTTGGGG | 74 | 74 |
| *Rdr6q-2r* | *Rdr6* – no exons | CCTTGAAGCTGTTGATGTGCC | 74 | 74 |
| *Rgd1q-1f* | *Rgd1* – exon 5,6 | GCCACATCCTTGACTTGGCA | 119 | 319 |
| *Rgd1q-1r* | *Rgd1* – exon 5,6 | TGCAGGAAGAGCGCTCAAAG | 119 | 319 |
| *Rgd1q-2f* | *Rgd1* – exon 4,6 | CGCTCTTCCTGCAACAACTT | 130 | 378 |
| *Rgd1q-2r* | *Rgd1* – exon 4,6 | GAGAAGCACATGGAGTACGAGG | 130 | 378 |
| *Ago1e-1f* | *Ago1e* – exon 19,20 | TTCTACCTGTGCAGCCATGC | 103 | 172 |
| *Ago1e-1r* | *Ago1e* – exon 19,20 | GAGTTTGCAACCCATCAGCC | 103 | 172 |
| *Ago1e-1f* | *Ago1e* – exon 10,11 | GCCAAAAATTGGCCAGTGGA | 128 | 515 |
| *Ago1e-1r* | *Ago1e* – exon 10,11 | TCATGACAGAAACTCCCGGC | 128 | 515 |
